# Evolutionary fingerprints of EMT in pancreatic cancers

**DOI:** 10.1101/2023.09.18.558231

**Authors:** Luigi Perelli, Li Zhang, Sarah Mangiameli, Andrew J. C. Russell, Francesca Giannese, Fuduan Peng, Federica Carbone, Courtney Le, Hania Khan, Francesca Citron, Melinda Soeung, Truong Nguyen Anh Lam, Sebastian Lundgren, Cihui Zhu, Desiree Catania, Ningping Feng, Enrico Gurreri, Alessandro Sgambato, Giampaolo Tortora, Giulio F. Draetta, Giovanni Tonon, Andrew Futreal, Virginia Giuliani, Alessandro Carugo, Andrea Viale, Timothy P. Heffernan, Linghua Wang, Davide Cittaro, Fei Chen, Giannicola Genovese

**Affiliations:** Department of Genitourinary Medical Oncology, The University of Texas MD Anderson Cancer Center, 1515 Holcombe Boulevard, Houston, Texas, 77030 USA; Broad Institute of Harvard and MIT, Cambridge, MA, 02142, USA; Department of Stem Cell and Regenerative Biology, Harvard University, Cambridge, MA 02138, USA; Center for Omics Sciences, IRCCS San Raffaele Institute, Milano, Italy; Department of Genomic Medicine, The University of Texas MD Anderson Cancer Center, 1515 Holcombe Boulevard, Houston, Texas, 77030 USA; Nerviano Medical Sciences, NMS Group Spa, 20014, Nerviano, Milan, Italy; TRACTION platform, The University of Texas MD Anderson Cancer Center, 1515 Holcombe Boulevard, Houston, Texas, 77030 USA; Medical Oncology, A. Gemelli University Hospital Foundation IRCCS, Largo A. Gemelli 8, Rome, Italy; Department of Translational Medicine and Surgery, Università Cattolica del Sacro Cuore, 00168 Rome, Italy; Department of Biology, IRBM S.p.A., Via Pontina Km 30 600, Pomezia, Rome 00071, Italy; The University of Texas MD Anderson Cancer Center UTHealth Houston Graduate School of Biomedical Sciences (GSBS), Houston, Texas, 77030 USA; David H. Koch Center for Applied Research of Genitourinary Cancers, The University of Texas MD Anderson Cancer Center, 1515 Holcombe Boulevard, Houston, TX 77030, USA

## Abstract

Mesenchymal plasticity has been extensively described in advanced and metastatic epithelial cancers; however, its functional role in malignant progression, metastatic dissemination and therapy response is controversial. More importantly, the role of epithelial mesenchymal transition (EMT) and cell plasticity in tumor heterogeneity, clonal selection and clonal evolution is poorly understood. Functionally, our work clarifies the contribution of EMT to malignant progression and metastasis in pancreatic cancer. We leveraged *ad hoc* somatic mosaic genome engineering, lineage tracing and ablation technologies and dynamic genetic reporters to trace and ablate tumor-specific lineages along the phenotypic spectrum of epithelial to mesenchymal plasticity. The experimental evidences clarify the essential contribution of mesenchymal lineages to pancreatic cancer evolution and metastatic dissemination. Spatial genomic analysis combined with single cell transcriptomic and epigenomic profiling of epithelial and mesenchymal lineages reveals that EMT promotes with the emergence of chromosomal instability (CIN). Specifically tumor lineages with mesenchymal features display highly conserved patterns of genomic evolution including complex structural genomic rearrangements and chromotriptic events. Genetic ablation of mesenchymal lineages robustly abolished these mutational processes and evolutionary patterns, as confirmed by cross species analysis of pancreatic and other human epithelial cancers. Mechanistically, we discovered that malignant cells with mesenchymal features display increased chromatin accessibility, particularly in the pericentromeric and centromeric regions, which in turn results in delayed mitosis and catastrophic cell division. Therefore, EMT favors the emergence of high-fitness tumor cells, strongly supporting the concept of a cell-state, lineage-restricted patterns of evolution, where cancer cell sub-clonal speciation is propagated to progenies only through restricted functional compartments. Restraining those evolutionary routes through genetic ablation of clones capable of mesenchymal plasticity and extinction of the derived lineages completely abrogates the malignant potential of one of the most aggressive form of human cancer.

## Main

Neoplastic cells display virtually unlimited evolutionary potential, which sustains tumor heterogeneity and ultimately drives lethal cancers^1–4^. This intrinsic trait of the tumor ecosystem is responsible for unlocking a multiplicity of distinct cell states, a hallmark of cancer defined as phenotypic plasticity^5^. Nevertheless, acquisition of phenotypic plasticity and its contribution to tumor progression and patients’ outcome remain poorly understood, particularly in regards of its role in metastatic dissemination, adaptation to therapy and evolutionary selection of aggressive clones^6–8^. As a paradigm of phenotypic plasticity, Epithelial-to-Mesenchymal Transition (EMT), a trans-differentiation program resulting in a continuum of cell-states, allowing the switch from epithelial to a spindle-like morphology, has been frequently described in advanced epithelial cancers and associated to dismal prognosis^9,10^. The mechanistic contribution of mesenchymal plasticity to cancer evolution, however, is still a subject of debate, as conflicting evidences from various research groups accumulated over the past two decades^11–15^.

To fill this knowledge gap, we set to clarify the functional role of EMT in cancer evolution focusing on pancreatic ductal adenocarcinoma (PDAC) as a model system where pathological and transcriptomic analysis of patients’ derived samples suggest that the emergence of mesenchymal features is associated with disease progression and poor survival^16–18^. In order to study the contribution of mesenchymal plasticity to cancer evolution we established somatic mosaic genetically engineered mouse model (SM-GEMM) of PDAC leveraging a dynamic lineage tracing and a lineage ablation systems allowing to trace and selectively remove the tumor lineages derived from cells undergoing EMT without perturbing the tumor microenvironment, which is a substantial tecnological advancement compared to previous *in vivo* technologies^11,19^. Our findings provide compelling evidence regarding the functional role of EMT as a crucial phenotypic program for malignant transformation and for the expansion of aggressive tumor subpopulations. Detailed annotation of genome-epigenome-phenome associations of SM-GEMM through bulk genomic, single cell and spatial sequencing technologies reveals that EMT is necessary to unlock the evolutionary potential of epithelial tumors towards aggressive clinical behaviour. Specifically, cross-species analysis of genomic signatures suggests that populations undergoing EMT are prone to acquire massive genomic instability characterized, in particular, by complex structural rearrangements and chromotripsis. Our work proves that highly plastic maligant subpopulations contribute to the generation of specific evolutionary routes and display unique patterns of structural genomic signatures, suggesting that diverse mechanisms of selection are at play in phenotypically distinct cancer cell populations.

### EMT is indispensable for malignant progression of pancreatic tumors

To study the role of mesenchymal plasticity in PDAC evolution we developed multiple reporter systems that allow for the tissue specific tracing of mesenchymal lineages (defined as malignant clones derived from an EMT-proficient ancestor). We engineered the mouse *Vimentin* locus for the conditional expression of a synthetic cassette harboring a H2B-GFP and a FLPO recombinase that is actively transcribed upon CRE-mediated recombination to ensure tissue and tumor cell specificity, excluding the stromal cellular compartment (*Vim^FLEX(SA–H2B–eGFP–T2A–FlpO–WPRE–pA)^*, hereafter *Vim^Flpo^*) (**Extended Fig. 1a**). Preliminary characterization of the SM-GEMM confirmed expected patterns of tissue expression and no overt phenotypic effect in both heterozygous and homozygous littermates (**Extended Fig. 1 b-c**). The reporter strain was further crossed into a somatic mosaic model of PDAC (R26^FSF-LSL-TdT^-H11^LSL-spCas^^9^-KRas^LSL-G12D^) where tumorigenesis is initiated through orthotopic delivery of Adeno-Associated Viral (AAV) particles carrying sgRNAs targeting the most common PDAC tumor suppressor genes, *Trp53* and *Cdkn2a/b,* along with a CRE-recombinase driving the activation of the conditional Cas9, the *KRas^G12D^* oncogene and the lineage tracing reporter in the edited pancreatic epithelial cells (hereafter **PCΦ**, please, see **Methods** for extended nomenclature). This model system allows for 3 possible experimental outputs: mesenchymal lineages (MS-S: TdT+/H2B-GFP^HIGH^), epithelial lineages (EPI-S: TdT**-**/H2B-GFP-) and mesenchymal competent lineages (MS-C: TdT+/H2B-GFP^LOW^) (**Figure 1a-b, Extended Fig. 1d**). Strikingly, gross and histopathological analysis of PCΦ models reveal that, advanced tumors and metastatic lesions were universally established by MS-S and MS-C lineages, while EPI-S cells were only detected in early neoplastic lesions (**Figure 1 c-g, Extended Fig. 2 a**). Histopathological, immunophenotypic and lineage tracing analysis of EMT in the PCΦ model also demonstrate that metastatic sites are almost universally established from MS-S and MS-C lineages and their growth is sustained by cells in an active mesenchymal state (TdT+/H2B- GFP^HIGH^) when compared to primary tumor sites (**Figure 1 h**). These evidences suggest that mesenchymal plasticity emerges very early during tumorigenesis, thus selecting for malignant populations with high fitness and metastatic competency. Data where further confirmed in orthotopic transplantation studies of TdT+ and TdT-organoid-derived cell populations confirming that TdT+ cells display aggressive clinical behavior, tendency to sarcomatoid transdifferentiation and metastatic spread (**Extended Fig. 2 b-f**). Remarkably TdT-transplants, while proving tumorigenic, resulted invariably in TdT+ lesions; this suggests that the activation of the EMT program is indispensable for lineages endowed with malignant potential, and that tumoral epithelial cells are highly meta-stable, thus prone to mesenchymal reprogramming. To functionally detail the contribution of EMT to pancreatic tumorigenesis, we engineered a lineage ablation system leveraging a DTA suicide cassette that selective kills malignant epithelial cells as they acquire mesenchymal features during tumorigenesis (**PCΨ, Methods**) (**Figure 1 i**). Strikingly, in line with the lineage tracing experiments, complete ablation of mesenchymal lineages strongly impacted tumor incidence, survival and metastatic burden resulting in virtually complete abrogation of malignant progression, with the emergence of low-grade cystic lesion lacking invasive potential and characterized by low proliferative index in both primary SM-GEMM and in orthotopic transplants (**Figure 1 j-o, Extended Fig. 2 g-j**). These evidences demonstrate that the ability of malignat cells to undergo EMT is dispensable for tumor initiation, but essential for the emergence and expansion of clones with highly proliferative, invasive and metastatic potential.

**Figure 1.**
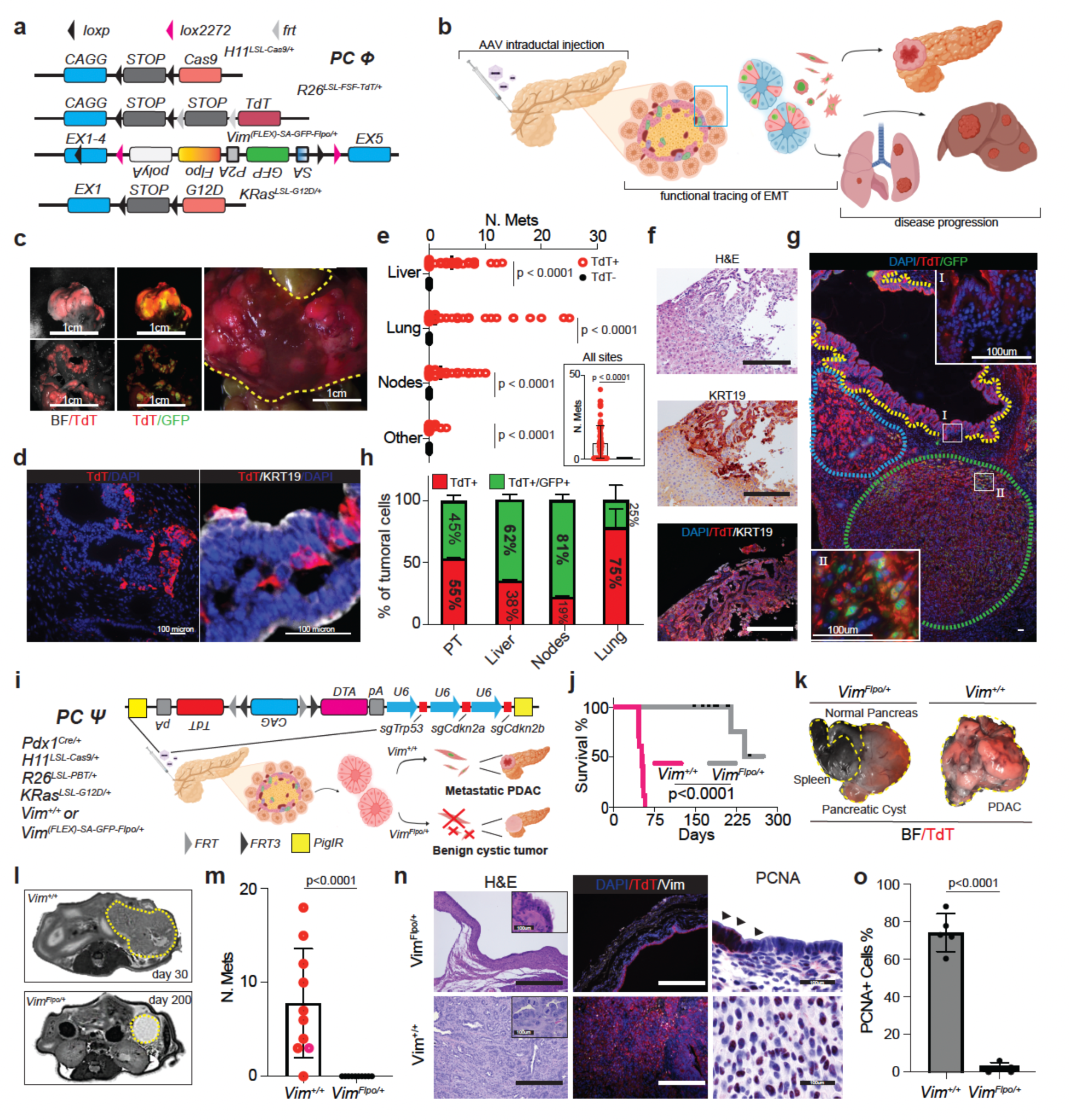
EMT is required for malignant progression of pancreatic tumors. **a-b)** Schematics of the alleles combination (**a**) generating a lineage tracing SM-GEMM of EMT in pancreatic cancer, initiated by the *KRas^G12D^* oncogene and different combinations of oncosuppressor knockouts achieved by CRISPR/Cas9 engineering (**b**). **c)** Macroscopic examination of a representative case of PCΦ SM-GEMM showing pervasive presence of TdT positivity at both primary and metastatic sites. Picture is representative of N = 76 mice. **d)** Histopathological characterization of a low grade pre-neoplastic lesion in the PCΦ SM-GEMM showing a mix of TdT+ cells in the context of KRT19+ PanIn lesions. **e)** Dot plot with individual values displaying total number of TdT+ and TdT-metastasis for the PCΦ model among four different metastatic sites and all metastasis combined. N = 76/mice included in the analysis. **f-g)** representative histopathological pictures showing a pervasive presence of TdT+ cells in a liver metastatic site **(f)** and a primary tumor site **(g)** both in low grade (I) and high grade (II) lesions. **h)** histopathological quantification of TdT+/GFP- and TdT+/GFP+ cells among primary tumor (PT) and three different metastatic sites. N = 25 lesions, 5 lesions/mouse per each anatomical site. **i)** Experimental schematic of the generation of the PCΨ model allowing for the conditional ablation of tumor cells expressing the synthetic *Vim^Flpo^* cassette. **j-k)** Survival curves of PCΨ *Vim^+/+^* and *Vim^Flpo/+^* mice; censored events are determined by mice sacrificed without evidence of tumor **(j)**. Macroscopic pictures of PCΨ *Vim^+/+^* and *Vim^Flpo/+^* mice-derived lesions (BF: brightfield and TdT); the cystic nature of *Vim^Flpo/+^*tumor is clearly visible without signs of infiltration of spleen and adjacent organsa. *Vim^+/+^*: N = 10; *Vim^Flpo/+^*: N = 10. Data are combined from two different experiments. Censored mice did not develop tumors. **l)** Representative MRI axial sections of PCΨ *Vim^+/+^* and *Vim^Flpo/+^* mice clearly displaying a fluid rich lesion in the *Vim^Flpo/+^* group in T2 scan. **m)** Dot plot displaying individual number of metastasis per each PCΨ *Vim^+/+^* and *Vim^Flpo/+^* mouse. *Vim^+/+^*: N = 10; *Vim^Flpo/+^*: N = 10. Data are combined from two different experiments. **n-o)** histopathological analysis of PCΨ *Vim^+/+^* and *Vim^Flpo/+^* mice-derived tumoral lesions evidencing the cystic appearance of *Vim^Flpo/+^*-derived tumors, characterized by low proliferation index, as determined by PCNA quantification **(o)**. N = 5 lesions per group. P values are calculated as follows: **j)** Logrank Mentel-Cox test; **e,m,o)** two-sided t-student test.

### Cells with mesenchymal features sustain tumor growth in advanced pancreatic cancer

We further tested whether cells with mesenchymal features are required to actively sustain the growth of advanced tumors. To this aim, we generated a second SM-GEMM by knocking-in a synthetic cassette harboring a EGFP and a HSV-TK (*Vim^FLEX(SA-eGFP-T2A-HSV-TK—WPRE-pA)^,* hereafter *Vim^TK^*) that is primed for transcription upon CRE-mediated recombination in the pancreatic epithelial compartment to ensure tumor specific ablation of mesenchymal lineages upon ganciclovir (GCV) administration. Specifically, the system allows for the tissue and time restricted ablation of malignant cells that are actively in a mesenchymal state while sparing the stromal and the epithelial compartments of the tumor (**PC**Ω) (**Figure 2 a, Extended Fig. 3 a-c**). This experimental model allowed us to distinguish 2 functional malignant subpopulations: one in an active mesenchymal state, TdT+/GFP^HIGH^ (MS-L), one lacking mesenchymal features TdT+/GFP^LOW^ (EPI-L). Indeed, GCV treatment of pancreatic tumors strongly inhibits tumor growth at primary and metastatic sites in both the short and long term experimental arms (21 and 90 days of treatment respectively). These effects are completely rescued after GCV pulse-release suggesting that the malignant epithelial compartment might function as a reservoir for the expansive mesenchymal compartment (**Figure 2 b-f**). Histopathological characterization of PCΩ tumors at the end of the treatment shows a significant depletion of malignant cells with mesenchymal features in the treated group, resulting predominantly in well differentiated epithelial structures (**Extended Fig. 3 d-e**). These evidences led us to the hypothesis that cells in active EMT are critical contributors of tumor growth, sustaining the homeostasis of the proliferative compartment. To further corroborate this hypothesis, we generated tumor explants from PCΩ mice to establish 3D cultures (**Extended Fig. 4 a**); in these experimental conditions GCV treatment resulted in a significant reduction in dimensions, without any relevant effect on clonogenic potential upon pulse-release (**Extended Fig. 4 b-c**). Furthermore, transplantation studies in orthotopic and heterotopic settings demonstrate that MS-L populations engraft and promote metastatic seeding more efficiently when compared to EPI-L as previously shown by our group ^20^(P<0.0001)(**Figure 2 g-j, Extended Fig. 4 d-n**).

**Figure 2.**
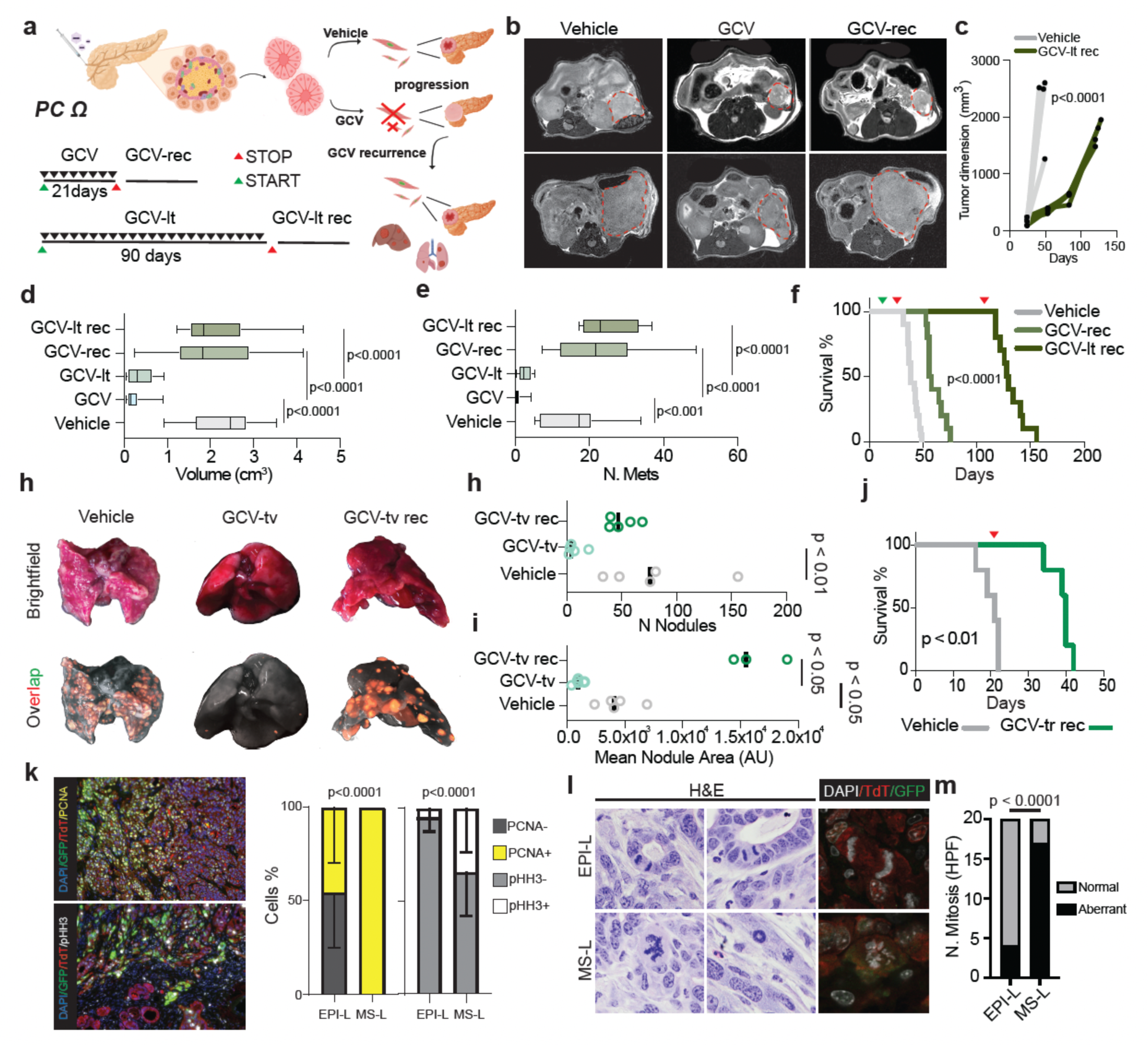
Tumor cells with mesenchymal features fuel tumor growth. **a)** Experimental schematic representing the PCΩ model and treatment schedule with GCV short term (21 days) and GCV long term (90 days) and respective recurrences at the end of treatment (GCV-rec and GCV-lt rec respectively). **b)** Representative T2 weighted axial MRI scans of PCΩ mice either treated either with Vehicle or GCV at morbidity (Vehicle and GCV-rec) and at the end of the treatment (GCV). **c)** Quantification of tumor volume via MRI imaging for the Vehicle and GCV-lt rec groups. N = 4 mice per group. **d-e)** Box plot showing quantification of tumor volume and number of metastases among the five previously described experimental arms. Vehicle: N = 20, GCV: N = 10, GCV-lt: N = 10; GCV-rec: N = 10; GCV-lt rec: N = 10. Data are extrapolated from two different experiments. **f)** Survival curves of Vehicle, GCV-rec and GCV-lt rec displaying a significant benefit in terms of survival with continuous pharmacological pressure. Vehicle: N = 20; GCV-rec: N = 10. **g)** Representative macroscopic pictures of lungs injected in the lateral tail vein with tumor-derived spheres and randomized to Vehicle or GCV (Vehicle: collection at morbidity; GCV-tv: collection at the end of treatment; GCV-tv rec collection at morbidity). Overlap: Brightfield, TdT and GFP. **h-i)** Quantification of number of nodules **(h)** and mean nodule area **(i)** displaying a significant reduction of number of nodules and mean nodule area in the GCV-tv group. N = 5 mice per group. **j)** Survival curves showing significant benefit of GCV treatment in transplantation studies. **k)** Histopatological analysis of proliferation potential of EPI-L (TdT+/GFP-) and MS-L (TdT+/GFP+) cells with representative pictures (left) and quantification (right) in primary tumors derived from the PCΩ SM-GEMM. N = 25 lesions from 5 different mice per group. **l-m)** representative images **(l)** and quantification **(m)** of aberrant mitotic (chromosome laggings, multipolar spindles and chromosomal bridges were included) figures in the two subpopulations. N = 5 different tumors. P values are calculated as follows: c,k) two-way ANOVA; **d,e,h,I)** two-sided student t-test; **f,j)** Logrank Mentel-Cox test; **m)** two-sided chi-square test.

Histopathological and cytological analysis of PCΩ pancreatic tumors reveals that malignant cells with mesenchymal features MS-L have higher proliferation index (P value < 0.0001) and a significant increase in the number of aberrant mitosis when compared to EPI-L cells, as assessed by PCNA and pHH3 stainings and through confocal microscopy of mitotic cells. These evidences altogether suggest that mesenchymal plasticity might increase the chance for mitotic errors (**Figure 2 k-m**). Our data collectively provide a counterintuitive model to the established role of EMT during malignant progression^10,21^, where mesenchymal plasticity is required for tumor growth and clonal expansion as demonstrated by GCV pulse-and-release experiments and transplantation studies.

### EMT promotes genomic instability and acquisition of catastrophic structural rearrangements

The functional studies described above point towards a potential role of mesenchymal plasticity as a functional driver of genomic instability in PDAC. To test this hypothesis, we performed Slide-DNA-seq^22^ on 3 independent tumors established from the PCΦ model 3 weeks after AAV injection in order to capture a spectrum of pre-neoplastic lesions and early maligant nodules. Strikingly, geospatial characterization of the genomic landscape of early tumors consistently showed the emergence of three clusters where the one overlapping with areas of TdT+ malignant mesenchymal cells uniformly displays features of aneuploidy including whole genome duplication (WGD) and chromosome level alterations including gains of 6 and 11, loss of 4 and 15, overall suggestive of ongoing chromosomal instability (CIN) (**Figure 3 a-c, Extended Fig. 5 a-d)**. These data prompted us to investigate wheter the genetic ablation of mesenchymal lineages restrains the evolutionary patterns of PDAC and the emergence of CIN with aberrant karyotypes; thus, we performed Whole Genome Sequencing (WGS) of malignant sub-populations isolated from tumors established from PCΨ and PCΩ SM-GEMM, where ablation of mesenchymal lineages is performed at different time points of tumor progression (collectively, **EMT-off** and **EMT-on**, please see **Methods**) aiming to inform on the impact of EMT-proficient lineages on cancer evolution.

**Figure 3.**
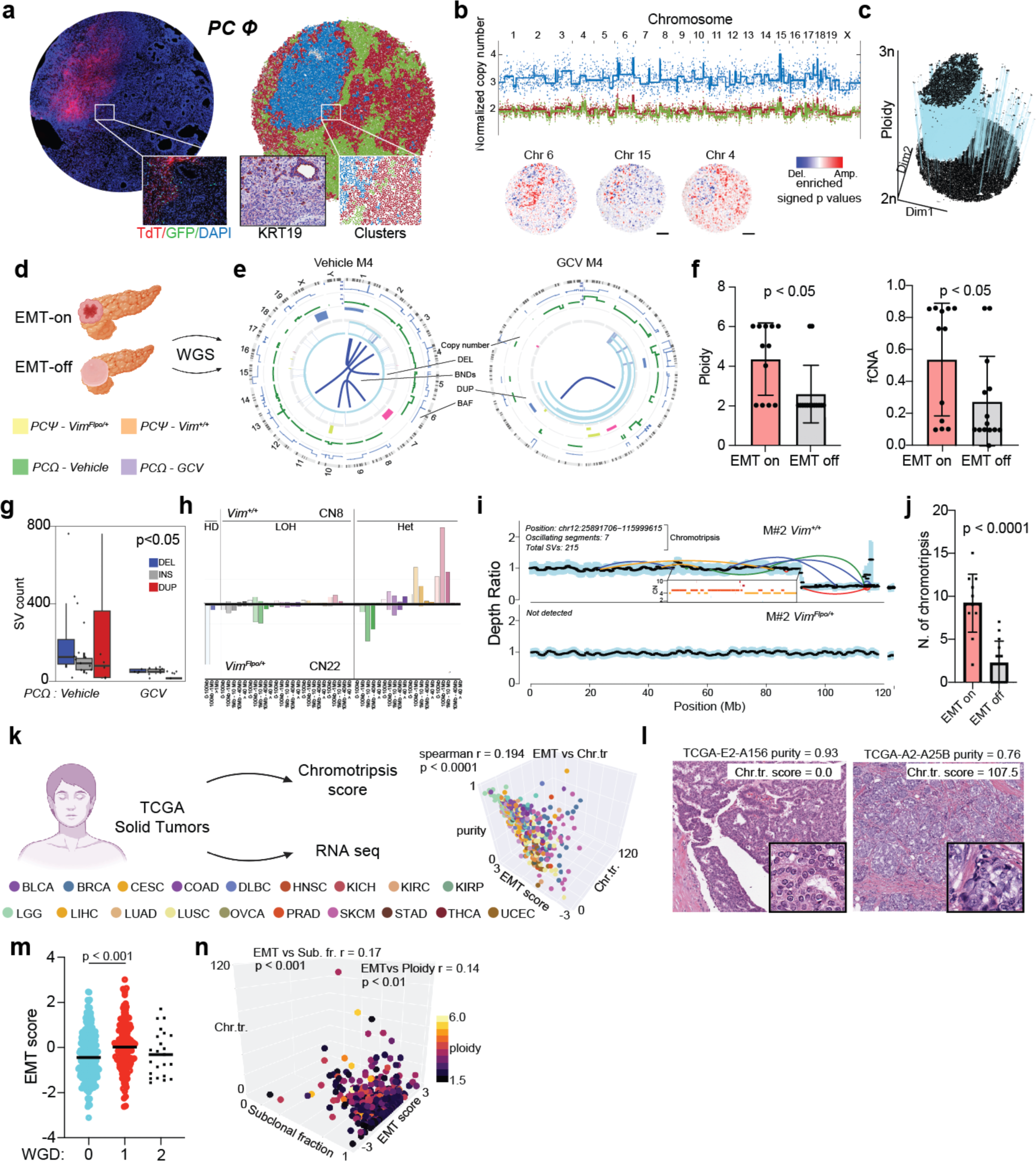
EMT promotes genomic instability and acquisition of events of chromotripsis. **a)** Slide-DNA-seq of a PCΦ SM-GEMM 3 weeks after AAV transduction showing TdT+ and TdT-regions of pancreatic tumor tissue, differentially clustering according to genomic analysis. The magnification evidences a TdT-/KRT19+ neoplastic lesion clustering differently from TdT+/KRT19+ cells. **b)** Normalized copy number scatter plot showing differential copy number aberrations in the three clusters (particularly evident gain of 6 and 15 and minor loss of chromosome 4, **b**). **c)** Three-dimensional plot showing geographic distribution of ploidy as calculated from Slide-DNA-seq. **d-e)** Experimental schematic and circos plot of two exemplary cases of PCΩ models clearly displaying enrichment of SVs and copy number events in the EMT-on sample. **f)** Summary plots showing individual values and mean of ploidy and fraction of copy number altered (fCNA) for EMT on and EMT off groups. **g)** Box and whiskers with data points showing number and type of SV enriched in the untreated mice. N = 7 mice per group. **h)** Copy number signatures for the EMT-on group (top) and EMT-off group (bottom) showing the prevalence of CN8 in the EMT-on group. i-j) Representative scatter plots and quantification of events of chromotripsis in EMT-on and EMT-off groups showing an increment of number of chromotripsis in the EMT-on group; a clear pattern of chromotripsis can be appreciated in the magnified box. **k)** Schematic and three-dimensional dot plot showing bioinformatic analysis correlating EMT score, Chromotripsis score (Chr.tr.) and purity from the TCGA cohort. **l)** Representative histopathological pictures of two breast cancer samples with low and high Chr.tr. Score and similar purity, where de-differentiation features can be appreciated with high levels of chromotripsis. **m)** Dot plot showing values of EMT score per each sample among WGD status (0: no WGD; 1: one WGD event; 2: two WDG events). **n)** Three-dimensional dot plot showing correlations between EMT score, estimated ploidy and tumor subclonal fraction (Sub. fr.). P values are calculated as follows: **g)** two-way ANOVA; **j)** two-sided student t-test; **k,o)** spearman correlation; **m)** two-sided Mann-Whitney.

Strikingly, ablation of EMT lineages results in relatively flat genomic landscapes, while the spontaneous expansion of malignant sub-populations with mesenchymal plasticity results into genomic features of aggressive and advanced PDAC^23^, specifically frequent copy number aberrations (CNAs), WGD and a significant increase in the number of complex structural variants (SVs) including deletions, duplications and insertions (P < 0.05). Such phenomena were evident collectively in both models, although more prominently in the PCΨ system, where mesenchymal lineages are genetically removed at the early stages of tumorigenesis, suggesting that context-dependent evolutionary patterns are highly specific for mesenchymal lineages (**Figure 3 d-g, Extended Fig. 6 a**). Therefore, we interrogate EMT-off and EMT-on tumors on different genetic processes driving progression through the analysis of copy number signatures demonstrating that EMT-on tumors are enriched for signatures related to chromotriptic events (CN8)^24^ as compared to EMT-off lesions, where these lineages are genetically depleted (**Figure 3 h**). Supporting these findings, we recovered the number of chromotriptic events per sample using ShatterSeek and by visual inspection of high-confidence chromotripsis events from each tumor (**Methods**)^25^. Strikingly, tumors with unrestrained potential for mesenchymal plasticity (EMT-on) display high burden of chromotripsis when compared to tumors where mesenchymal lineages are ablated (EMT-off, P < 0.0001) (**Figure 3 h-j, Extended Fig. 6 b-c**). Cross-species analysis of patients’ derived malignant lesions (TCGA) annotated for number of chromotriptic events and transcriptomic information further confirmed that epithelial tumors with transcriptomic features of mesenchymal reprogramming are prone to accumulate chromotriptic events, WGD and subclonal SVs, leading to overall poorer survival. These phenomena are tissue agnostic, as being observed across anatomical disease sites and histological sub-types, strongly supporting the hypothesis that mesenchymal plasticity is permissive for the emergence and selection of complex structural rearrangements and karyotypes (**Figure 3 g-o**).

To understand the functional impact of chromotriptic events and other structural genomic rearrangments related to EMT during pancreatic cancers progression, we examined chromosomes affected by chromotripsis at high frequency and SVs in the EMT-on group (**Figure 4 a-b**). In particular, chromosomes 4 and 6 chromotripsis were always present in this cohort. Surpisingly, tumors belonging to the EMT-on group were enriched with SVs affecting the cluster of Interferon ligands genes on chromosome 4C4 (P < 0.05) mostly leading to intragenic inversions and deletions through intra-exonic breakpoints, affecting genes integrity (**Figure 4 c-d**). Furthermore, visualization of the chromotripsis regions on chromosomes 6 revealed copy number amplification of oncogenic drivers, particularly gain of the *KRas* oncogene (**Figure 4 e-f**). Allele analysis of the *KRas* gene locus demonstrated a strong enrichment of *KRas* allelic imbalance in the EMT-on group. These data are strongly suggestive of an active role of EMT in shaping pancreatic cancer genomic evolution through the acquisition of key drivers events^26,27^.

**Figure 4.**
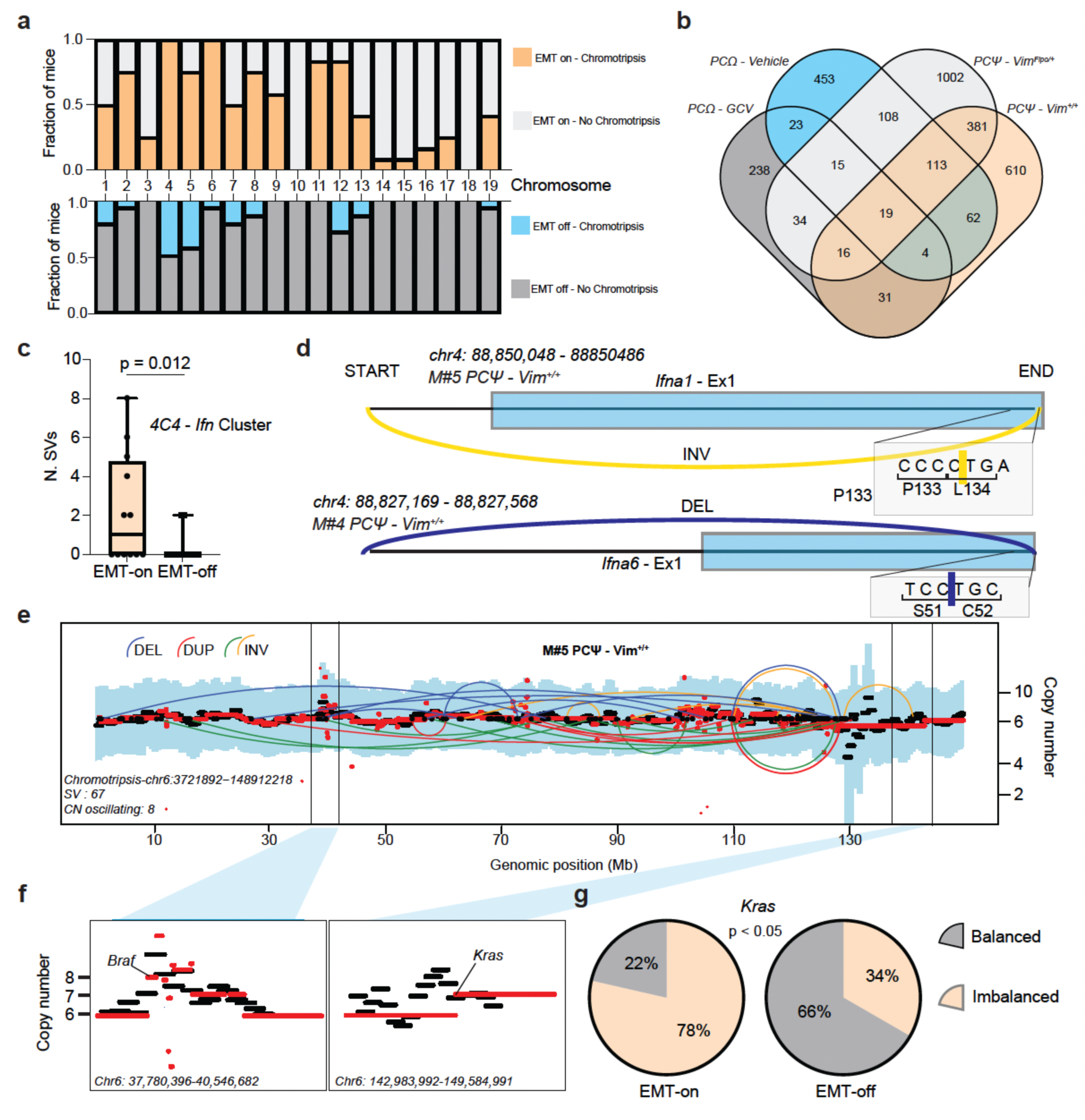
Acquisition of driver events during EMT in pancreatic cancer. **a)** Bar plots showing fraction of mice with chromosome-specific events of chromotripsis among the EMT-on and the EMT-off group. **b)** Venn diagramm showing numbers of private and shared genes affected by chromotripsis among different SM-GEMM as calculated by WGS analysis. **c)** Bar plot displaying number of SVs per each mouse with breakpoints on the 4C4 cluster of genes of Interferon ligands (*Ifn* Cluster). **d)** Representative inversion (top) and deletion (bottom) involving two genes of the *Ifn* Cluster. **e)** Representative case of chromotripsis of chromosome 6 in a mouse belonging to the EMT-on group. **f)** Zoom in on two genomic regions on chromosome 6 showing amplification of the *Braf* and *Kras* oncogenes. **g)** Pie charts showing percentages of mice with *Kras* imbalance among the EMT-on (left) and EMT-off (right) groups. P values are calculated as follows: **c)** two-sided student t-test; **g)** Fisher’s exact chi-test.

### Increased chromatin accessibility drives genomic instability and functional diversification of PDAC

To understand the impact of mesenchymal plasticity on subclonal speciation in PDAC and functional tumor heterogeneity, we performed single cell RNA and GET^28^ sequencing on short term cultures of sorted MS-L and EPI-L subpopulations isolated from the PCΩ SM-GEMM, providing informations on functional cell states, genomic landscape and chromatin dynamics associated with such functional states at the single cell level. Integration of epigenetic and transcriptomic profiles, confirmed the presence of 2 different states defined by the activation of EMT programs (**Extended Fig. 7 a-d**). Remarkably, MS-L cells display increased genomic instability, accumulation of genomic SVs and clonotypic diversity along the EMT continuum as displayed by integration of single cell RNA and GET sequencing data (P < 0.0001) (**Figure 5 a, Extended data Fig. 7 e-g, Methods**). Moreover, detailed inference of genomic complexity from scGET-seq data reveals an enrichment of SVs in chromatin regions with high accessibility, in particular centromeric and peri-centromeric chromosomal regions, overall characterized by a global increase in accessibility (**Figure 5 b-e**). These evidences suggest that alteration of pericentromeric chromatin compaction might result in delayed mitosis, lagging chromosomes and extranuclear chromatin sequestration which are hallmarks of CIN and permissive of catastrofic genomic events like chromotripsis (**Figure 5 f**)^29,30^.

**Figure 5.**
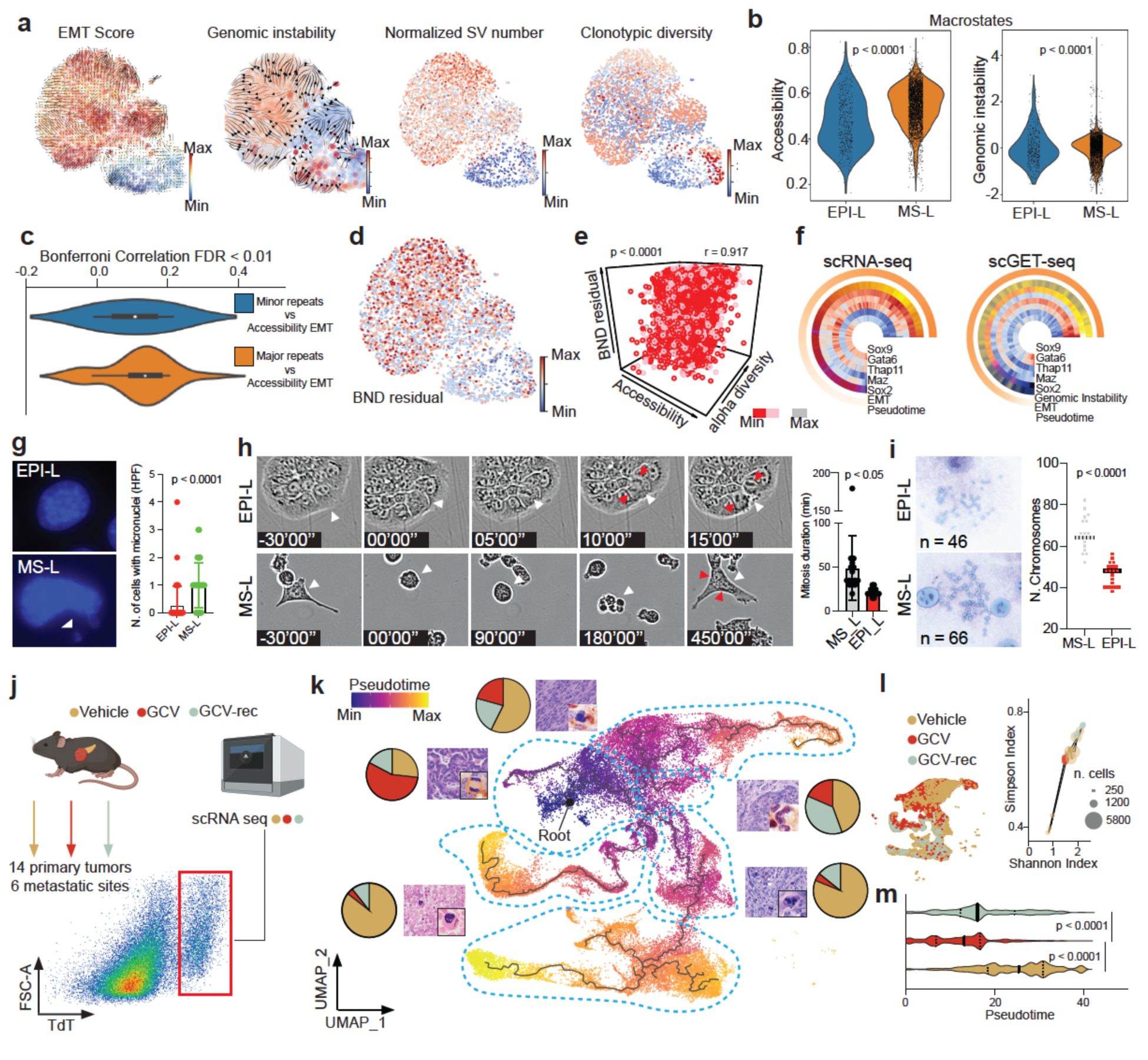
Chromatin accessibility drives genomic instability and functional heterogeneity during EMT. **a)** UMAP showing EMT Score, Genomic instability, Normalized SV number and clonotypic diversity as calculated by integration of scRNA and scGET seq data of isogenic EPI-L and MS-L subpopulations established from the same PCΩ tumor model. **b-c)** Violin plots showing scGET seq accessibility and genomic instability values **(b)** and single cell correlation values for co-occurrence of SV in Minor and Major repeats and EMT continuum **(c)**. **d-e)** UMAP showing number of SV-breakpoints (BND-residual) **(d)** and three-dimensional dot plot showing single cell distribution, and correlation, among Accessibility, BND-residual and alpha diversity in sc-GET seq data **(e)**. **f)** Pie chart showing correlations among EMT score, cellular differentiation (pseudotime) and genomic instability with top significant transcription factor in the dataset based on expression and activity as calculated from scRNA and GET seq data. **g)** Representative picture (left) and quantification (right) of presence and number of micronuclei/area across EPI-L and MS-L subpopulations. **h)** Time-lapse frames (left) and quantification (right) of mitotic time in EPI-L (top) and MS-L (bottom) subpopulations. The analysis demonstrates increased mitotic time in the MS-L population. **i)** Representative pictures and quantification of metaphasic spreads of EPI-L and MS-L subpopulations showing higher median number of chromosomes in MS-L subpopulation. **j-k)** Schematic and UMAP showing scRNA sequencing data processing of PCΩ tumors along pseudotime using as root of origin acinar cells. Micrographs display putative phenotypes associated with transcriptomic features. **l)** Dot plot showing increased Simpson Index and Shannon Index (**Methods**) values in samples belonging to the Vehicle and GCV-rec group. **m)** Violin plot showing increased single cells Pseudotime values in the Vehicle and GCV-rec group. P values are calculated as follows: **b,g,i,m)** paired student t-test; **e)** pearson correlation.

Leveraging the PCΩ *in vivo* and *in vitro* data, we demostate that MS-L subpopulations displays a significantly greater number of micronuclei, prolonged time of mitosis, increased number of chromosomes per cell, larger nuclear sizes and an increased activation of DNA damage pathway as assessed by γH2AX staining, all indicative of mitotic and nuclear abnormalities leading to complex structural chromosomal aberrations as previously described (**Figure 5 g-i, Extended Fig. 7 h-i**). We further speculated that increased chromatin accessibility and mitotic errors in cancer cells may support the emergence of clonal functional diversification in PDAC. Single cell RNA sequencing date were therefore obtained in the PCΩ model from 14 primary tumors and 6 metastatic sites and 3 experimental conditions Vehicle, GCV and recurrence after GCV pulse-chase (**Figure 5 j**). After sorting of TdT+ populations and computational exclusion of TdT-cells, a total number of 41130 high confidence tumor cells were retrieved from both primary and metastatic lesions Deconvolution of lineage trajectories of single cell RNA sequencing data demonstrate that PDAC display high level of intratumor heterogeneity which is mostly due to the contribution of cancer cells belonging to untreated (Vehicle) and recurrent (GCV-rec) tumors, while treated (GCV) tumors display low level of transcriptomic entropy, as calculated with two independent indeces (Shannon and Simpson, **Methods**) (**Figure 5 k-m**). Furthermore, we discover that transcriptional programs of cells undergoing EMT strongly correlate with transdifferentiation and metastatic processes, while anti-correlating with ductal phenotype (“Classical” score) (**Extended data Fig. 8 a-h**). Finally, computational inference of sub-clonal diversification of malignant cells shows that Vehicle treated tumors display a higher number of clonotypes when compared to GCV tumors, further providing evidence of subclonal speciation of cancer cells undergoing EMT as a determining factor of functional heterogeneity (**Extended Fig. 8 i-j**).

## Discussion

Our work clarify the role of cell plasticity as a complex biological program observed in several solid tumor types, where it has been generally described as a response to external cues or during natural evolution of malignant tumors. Specifically, EMT has been observed to be a key feature influencing the metastatic process^19^.

While the major paradigm suggests that malignant epithelial cells must activate EMT to become invasive and metastatic, favoring cell motility over proliferation potential, we demonstrate through selective tracing and ablation of cell state specific tumor lineages that the acquisition of mesenchymal competency is necessary for malignant progression and tumor growth, promoting the emergence of high-fitness clones characterized by complex patterns of aneuploidy. This is in line with numerous clinical and histopathological evidences showing that epithelial cancers, characterized by high grade histologies and/or sarcomatoid dedifferentiation, display increased mitotic rate, proliferative capacity and genomic instability^16,31,32^. Indeed, in the context of PDAC, basal-like or quasi-mesenchymal tumors have been associated with increased proliferation rate and aggressive biological behaviour ^17,33^.

The emergence of specific copy number signatures in a distinct cell-state specific malignant subpopulation suggests that diverse mutational processes are at play during cell plasticity dynamics, thus contributing to clonal speciation and tumor heterogeneity. It is reasonable to speculate that the poor progrosis universally associated with the emergence of mesenchymal features in solid tumors might be a consequence of the selection of phenomena of punctuated evolution followed by rapid clonal sweeps in specific transcriptomic/functional cell states^1,2^. This is strongly supported by our experimental findings, whereof the selective ablation of the mesenchymal lineages *in vivo* results in relatively flat genomes. Our data are recapitulated by genomic and transcriptomic patients’ datasets, where lesions characterized by transcriptomic features mesenchymal competency are pervasively accumulating chromotripsis and other complex structural aberrations. Remarkably, such events are observed with the highest frequency in primary sarcomas resulting from the malignant transformation of mesenchymal or mesenchymal progenitor cells ^25^. It would be of great interest to investigate whether the mesenchymal state (constitutive or acquired through the process of EMT) is universally more permissive or tolerogenic to complex and catastrophic structural rearrangments including chromothripsis.

Mechanistically, we discovered that structural genomic aberrations correlate with a “relaxed” and accessible chromatin state, particularly in the peri-centromeric and centrometic regions that significantly increase the chances of mitotic delays and chromosome lagging. Recent evidences, and our previous work, shed light on the critical role of chromatin remodeling during mitosis as a bookmark of lineage fidelity, in particular in the context of SWI/SNF deficient cancer cells^20,34^. Altogether, these data point towards an active role of cell plasticity and chromatin remodelling in the genomic evolution of aggressive neoplasms through acquisition of CIN, leading to functional diversification of cancer cells, an overall increase in tumor entropy.

In conclusion, our findings have important conceptual implications for the field of tumor evolution, particularly introducing the concept of cell state specific evolutionary patterns. Seminal work from Stratton and others has led to the identification of conserved signatures of mutational processes and structural variants in human cancers, informing on the effect of specific oncogenic insults on the genomic landscape of malignant tumors^25,35,36^. Here we highlight a novel level of complexity, where different functional subpopulations within a tumor can evolve according to diverse mutational processes that demonstrate a degree of cell state specificity in such processes. These data add another layer of complexity to the concept of intratumoral heterogeneity and tumor cell plasticity as a driver of genomic and functional heterogeneity in PDAC and potentially other aggressive solid tumors.

## Methods

### Genetic engineered mouse models

The Vim^FLEX(SA–H2B–eGFP–T2a–FlpO–WPRE–pA)^ strain was generated by genOway, targeting exons 2 to 4 of the mouse Vim locus; the design has been carried out to minimize the risk of interference of regulatory elements. The Vim^FLEX(SA-eGFP-T2A-HSV-TK-WPRE-pA)^ strain was generated by genOway, targeting exons 2 to 4 of the mouse Vim locus; the design has been carried out to minimize the risk of interference of regulatory elements. The H11^LSL-Cas^^9^ strain was generated by Dr. Monte M. Winslow and obtained through the Jackson Laboratory, stock No: 027632^37^. The Rosa26^LSL-TdT^ was generated in Dr. Hongkui Zeng’s laboratory and obtained through the Jackson Laboratory, stock No: 007908^38^. The Rosa26^FSF-LSL-TdT^ was generated in Hongkui Zeng’s lab and obtained through the Jackson Laboratory, stock No: 021875^39^. The Kras^LSL-G12D^ was generated in Dr. Tyler Jacks lab and obtained through Jackson Laboratory, stock No: 008179^40^. The Rosa26^LSL-PBT^ was kindly donated by dr. Rolan Rad^41^. The Pdx1^CRE^ strain was generated by Dr. David Tuveson and obtained through Jackson Laboratory, stock No: 014647^42^. The Ella^CRE^ was generated in Dr. Heiner Westphal’s laboratory and obtained through the Jackson Laboratory, stock No: 003724^43^. Strains were kept in a mixed C57BL/6 and 129Sv/Jae background. CB17SC-F SCID mice were purchased from Taconic. All animal studies and procedures were approved by the UTMDACC Institutional Animal Care and Use Committee. All experiments conformed to the relevant regulatory standards and were overseen by the institutional review board. No sex bias was introduced during the generation of experimental cohorts. Littermates of the same sex were assigned randomly to experimental arms.

### GEMM nomenclature

PCΦ: Rosa26FSF-LSL-TdT, H11LSL-spCas9, KRasLSL-G12D, VimFLEX(SA-H2B-eGFP-T2a-FlpO-WPRE-pA)_;_ PCΨ: Pdx1Cre, Rosa26LSL-PBT, H11LSL-spCas9, KRasLSL-G12D, VimFLEX(SA-H2B-eGFP-T2a-FlpO-WPRE-pA)_; PCΩ:_ Rosa26LSL-TdT, H11LSL-spCas9, KRasLSL-G12D, VimFLEX(SA-eGFP-T2A-HSV-TK-WPRE-pA). PCΦ, PCΨ and PCΩ models were tested for the following combination of AAV-CRE: KPCC (sgTrp53, sgCdkn2a and sgCdkn2b. EMT-on group is composed by PCΨ-*Vim*^+/+^ and PCΩ-Vehicle mice; EMT-off group is composed by PCΨ-*Vim*^Flpo/+^ and PCΩ-GCV mice.

### Animal procedures

*<u>Retrograde ductal injection:</u>* retrograde ductal injection was performed as previously described^37^. Mice were anesthetized using isofluorane (Henry Schein Animal Health). Analgesia was achieved with buprenorphine SR (0.1 mg/Kg BID) (Par Parmaceutical) via subcutaneous injection, and shaved skin was disinfected with 70% ethanol and betadine (Dynarex). Laparotomy was performed through a midline incision of 1cm starting from the xyphoid process. The appendix was located and pulled out of the abdomen along with about half of the total bowel mass, exposing the duodenum. The sphincter of Oddi was exposed and injection of the AAV or Ad particles was performed with a 30-gauge needle, inserted through the sphincter of Oddi into the common bile duct up to where the cystic duct converges. About 100µL of 10^10^ viral particles resuspended in 1X PBS was injected. Then, the needle was slowly removed 30 seconds after completion of the injection. Muscular/peritoneal planes were closed individually by absorbable sutures. The skin/subcutaneous planes were closed using metal clips. Mice were monitored daily for the first three days, and twice/week.

*<u>Orthotopic parenchymal pancreatic injection:</u>* tumor-explant-derived three-dimensional cultures and genetically engineered pancreatic spheroids were resuspended in single cell suspension using trypsin-based method at a concentration of 1000 cells/µL in 1X PBS. Mice were anesthetized using isofluorane (Henry Schein Animal Health). Shaved skin was disinfected with betadine and ethanol and 1-cm incisions were performed through the skin/subcutaneous and muscular/peritoneal layers. The spleen and tail of the pancreas were identified, exposed, and one injection was performed in the pancreatic tail and body. The muscular/peritoneal planes were closed using continuous absorbable sutures. The skin/subcutaneous planes were closed using interrupted absorbable sutures. Analgesia was achieved with buprenorphine (0.1 mg kg-1 BID).

*<u>Intrasplenic injections:</u>* for intrasplenic injections, tumor-explant-derived three-dimensional cultures were resuspended in single cell suspension using trypsin based method at a concentration of 1000 cells/µL in 1X PBS. Mice were anesthetized using isofluorane (Henry Schein Animal Health). Once the animals were under deep anaesthesia, the area of injection was disinfected with Betadine and 70% ethanol. The spleen was exposed through a small incision. Cells were injected into the spleen with a single injection using an insulin syringe. Cells were given 10 min to travel through the vasculature to the liver, after which the entire spleen was removed to prevent the formation of a large splenic tumour mass. To remove the spleen, a dissolvable 4-0 suture was tied snugly around the base of the spleen including the major splenic vasculature and the spleen was removed; to achieve hemostasis, a cauterized was used when needed. The muscular/peritoneal planes were closed using continuous absorbable sutures. The skin/subcutaneous planes were closed using interrupted absorbable sutures. Analgesia was achieved with buprenorphine (0.1 mg kg-1 BID).

*<u>Intravenous injection:</u>* tumor-explant-derived three-dimensional cultures were resuspended in single cell suspension using trypsin-based method at a concentration of 1000 cells/µL in 1X PBS. Mice were anesthetized using isofluorane (Henry Schein Animal Health). Cells were intravenously injected through the lateral vein. Mice we monitored three times per week until moribound.

*<u>Treatments:</u>* When average tumor size reached ∼500mm^3^ of volume (around day 21 according to MRI imaging) mice received daily intraperitoneal injections with 50mg/kg of ganciclovir (GCV, Invivogen). Treatment schedules are specified for each experiment.

*<u>Euthanasia, necropsy, and tissues collection:</u>* mice were euthanized by exposure to CO2 followed by cervical dislocation. A necropsy form was filled in with mouse information, tumor size and weight, infiltrated organ annotations, and metastasis number and location. Fluorescent and brightfield images were acquired with Leica. Euthanasia was performed with animals at clinical terminal disease and metastatic tumor burden.

### Recombinant DNA

Packages of one or more guide RNAs were designed and synthetized according to the following scheme: MluI restriction site-U6 promoter-gRNA1 sequence-gRNA scaffold-polyA-U6 promoter-gRNAn sequence-gRNA scaffold-polyA-KpnI restriction site. The synthetic sequence was assembled into the AAV:ITR-U6-sgRNA(backbone)-pCBh-Cre-WPRE-hGHpA-ITR vector (Addgene)^44^ into the MluI and KpnI restriction sites. The Ad-PB::reverse complement TdT-(FLEX-FRT)reverse complement CAGG-DTA-polyA-U6 promoter-gRNA1 sequence-gRNA scaffold-polyA-U6 promoter-gRNAn sequence-gRNA scaffold–polyA was constructed as follows: first the PB inverse repeats 5’-reverse complement TdT-(FLEX-FRT)reverse complement CAGG-DTA-polyA-MluI restriction site-KpnI restriction site-PB inverse repeats 3’ was synthetized and clone into the pUC57 into the EcorI and XbaI restriction sites; second the U6 promoter-gRNA1 sequence-gRNA scaffold-polyA-U6 promoter-gRNAn sequence-gRNA scaffold–polyA was derived from the previously described AAV:ITR-U6-sgRNA(backbone)-pCBh-Cre-WPRE-hGHpA-ITR vector and cloned into the pUC57-PB inverse repeats 5’-reverse complement TdT-(FLEX-FRT)reverse complement CAGG-DTA-polyA-MluI restriction site-KpnI restriction site-PB inverse repeats 3’ into the MluI and KpnI restriction sites; the final construct was assembled into the Adeno-X adenoviral system (Takara Bio) via In-Fusion cloning as described by manufacturer protocol.

### Virus production

Plasmid DNA preparations were generated using endotoxin-free MIDI kits (Qiagen). Large-scale AAV and Ad particle production was outsourced to Vector Biolab (10^13 IU/mL).

### Non-invasive imaging

A 7T Bruker Biospec (BrukerBioSpin), equipped with 35mm inner diameter volume coil and 12 cm inner-diameter gradients, was used for MRI imaging. A fast acquisition with relaxation enhancement sequence with 2,000/39 ms TR/TE, 256×192 matrix size, r156uM resolution, 0.75 mm slice thickness, 0.25 mm slice gap, 40 x 30 cm FOV, 101 kHz bandwidth, and 4 NEX was used for acquired in coronal and axial geometries a multi-slice T2-weighted images. To reduce respiratory motion, the axial scan sequences were respiratory gated. All animal imaging, preparation, and maintenance was carried out in accordance with MD Anderson’s Institutional Animal Care and Use Committee policies and procedures.

### Pancreatic spheroids

Spheroid cultures were maintained as previously described with few modifications^45^. Pancreata were isolated and ductal fragments were isolated by collagenase digestion (C9407, Sigma) for 30 min at 1 mg/ml. Fragments were seeded in growth factor-reduced Matrigel (Corning) and cultured in medium (DMEM/F12 supplemented with 1% penicillin/streptomycin, HEPES, GlutaMAX), with 2% B27 supplement (Gibco), recombinant mouse noggin (50ng/ml Preprotech), 10% Rspo1 (Millipore-Sigma, SCM104), EGF (50 ng/ml, Peprotech), FGF-10 (100 ng/ml, Preprotech), N-acetylcysteine (1.25 mM, Sigma), A8301 (5 µM, Tocris Bioscience) and primocine (0.1 mg/ml, Invivogen). After 2 weeks, pancreatic spheroids were cultured using DMEM/F12 supplemented with 1% P/S, 10% FBS. After 3 weeks, pancreatic spheroids were dissociated from Matrigel in ice cold PBS, collected and pelleted. Organoids were plated at high confluency in 96 wells with DMEM/F12 supplemented with 1% P/S, 10% FBS and incubated with AAV (10^7^ viral particles) for 8 hours at 37° C 5% CO2. Single cell suspensions were collected, pelleted and embedded in Matrigel or transplanted, as previously described.

### Tumor cell isolation, sorting and culture

*Ex vivo* cultures from primary tumor explants were generated by mechanical dissociation and incubation for 1 hour at 37°C with a solution of collagenase IV/dispase (2 mg/ml) (Invitrogen), resuspended in DMEM (Lonza) and filtered. Cells derived from tumor dissociation and digestion were resuspended as single cell suspensions in 1X PBS and 5% FBS. Single cell suspensions were processed through fluorescent-activated cell sorting (FACS) using tdTomato fluorescence as a reporter for tumor cells. Sorted tdTomato+ cells were centrifuged at 200g per 5 minutes and resuspended in the appropriate medium.

*<u>Sphere formation assay and treatment:</u>* For sphere forming assay, 25-50k cells were carefully resuspended as single cell suspension in 300 mL of Stem Cell medium (SCM). Resuspended cells were then gently mixed with 1.5 mL of reconstituted MethoCult (StemCell Technologies, cat. n. M3134) and aliquot in a single well of an ultra-low attachment 6-well plate (Corning Costar, cat. n. 3471). Cells were kept at 37C and 5% CO2 for 5-10 days to allow sphere formation from single cells. Images of spheres were captured using Nikon microscope, sphere number and size were measured using NiS Element software (Nikon). Second generation spheres were plated as described above and treated with 10 microM GCV.

SCM preparation: MEBM (Lonza, cat. n. CC-3153) was supplemented with 1% Penicillin/Streptomycin (100 U/ml), 2 mM L-Glutammine (Sigma-Millipore, cat.n. 59202C), 0.5 mM Hydrocortisone (Sigma-Millipore, cat.n. H0135), 1X serum-free B27 Supplement (Gibco, cat.n. 17504044), 20 ng/mL mouse recombinant EGF (Peprotech, cat.n 315-09), 20 ng/mL human recombinant FGF (Peprotech, cat. n. 100-18B), 1X ITS liquid media supplement (Sigma-Millipore, cat.n. I3146), 100 mM 2-mercaptoethanol (Sigma-Millipore, cat.n. M3148). MethoCult reconstitution: 40 mL of Methocult were diluted to a final volume of 100 mL using MEMB and supplemented as described above for SCM.

*<u>EPI-L and MS-L isolation:</u>* sorted cells were cultured as two-dimensional cultures in 0.1% gelatin coated plates in DMEM High Glucose, 10% FBS, 1% P/S. After attachment and stabilization, cell cultures were either treated with 10µM GCV or with an equal volume of PBS in DMEM High Glucose, 10% FBS, 1% P/S to isolate EPI-L and MS-L populations respectively. Medium was changed every 24 hours for both EPI-L and MS-L cell cultures. Five passages after treatment, cells were subjected to downstream analysis.

### Single-guide RNA design

Single-guide RNAs (sgRNAs) sequences used in this study were obtained through Genescript and validated elsewhere(cit., link).

Single-guide RNA sequences: *Trp53*: CATAAGGTACCACCACGCTG, *Cdkn2a*: GTGCGATATTTGCGTTCCGC, *Cdkn2b*: GGCGCCTCCCGAAGCGGTTC.

### Staining

Immunohistochemistry (IHC) and immunofluorescence were performed as previously described^46^. Antibodies list: RFP (Thermo Fisher, cat. #MA5-15257), GFP (Abcam, cat. #13970), Vimentin (Abcam, cat. #ab8978), .

Multispectral imaging using the Vectra. Microwave treatment (MWT) was applied to perform antigen retrieval, quench endogenous peroxidases, and remove antibodies from earlier staining procedures. Akoya Biosciences AR6 antigen retrieval buffer (pH 6) was used for vimentin and RFP staining. The slides were stained with primary antibodies against RFP, GFP, and Vimentin, corresponding HRP conjugated secondary antibodies, and subsequently TSA dyes to generate Opal signal (vimentin, Opal 570; RFP, Opal 620; and Pax8, Opal 690). The slides were scanned with the Vectra 3 image scanning system (Caliper Life Sciences), and signals were unmixed and reconstructed into a composite image with Vectra inForm software 2.4.8.

### Metaphase spread and chromosome count

Giemsa staining on metaphasic spread was obtained as previously described with few modifications. Cultures were treated with 100 ng/ml nocodazole for 8 hours overnight, collected by trypsinization, resuspended in 0.2% (w/v) KCl and 0.2% (w/v) trisodium citrate hypotonic buffer at room temperature (20–22°C) for 5 to 10 min, and cytocentrifuged onto SuperFrost Plus glass slides (MenzelGlaser) at 450g for 10 min in a Shandon Cytospin 4. Slides were fixed at room temperature for 10 min in 1× PBS with 4% (v/v) formaldehyde, permeabilized for 10 min at room temperature in KCM buffer (120 mM KCl, 20 mM NaCl, 10 mM Tris (pH 7.5) and 0.1% (v/v) Triton X-100), and blocked with 5% goat serum PBS1X 0.1%Triton 100x BSA 3% for 30 min at room temperature. Slides were incubated with Giemsa solution for 5 minutes at room temperature.

### Next-generation sequencing of murine DNA

Exome libraries and whole genome-libraries were prepared and sequenced using a modified protocol^47^. Modifications to the protocol for murine exome sequencing were the use of 1000ng of treated gDNA, performing only 6 cycles of PCR amplification, and usage of the Agilent SureSelectXT Mouse All Exon Kit for exon target capture. For murine whole-genome sequencing, after adapter ligation, libraries were only amplified by 2 cycles of PCR. Equimolar quantities of the whole-genome indexed libraries were multiplexed, with 18 libraries per pool. Results from 13 of the 18 libraries were used in our analysis. All pooled libraries were sequenced on an Illumina NovaSeq6000 using the 150 bp paired-end format.

### Bioinformatic processing of high-throughput sequencing data

The bioinformatic processing pipeline of raw whole-exome and whole-genome (WGS) high-throughput sequencing data was adapted for murine data^48^. Reads were aligned to the mouse genome reference (mm10) using Burrows-Wheeler Aligner (BWA) with a seed length of 40 and a maximum edit distance of 3 (allowing for distance % 2 in the seed). BAM files were further processed according to GATK Best Practices, including removal of duplicate reads, realignment around indels, and base recalibration.

### Analysis of sgRNA performanc*e*

Expected cut sites of sgRNAs were analyzed using CRISPResso2. BAM files were first filtered with SAMtools to contain reads spanning a 50-bp region centered around the expected sgRNA cut site and passed to CRISPResso2 in “CRISPRessoWGS” mode. The allele frequency of each base position around the cut site window was extracted from the CRISPResso2 results^49^.

### Identification and characterization of somatic mutations

Somatic mutations were detected from murine tumor samples using a combination of MuTect v2^50^ to call somatic single-nucleotide variants (SNVs). Tumor samples from WGS were compared to their respective matched healthy tissue. All mutations were also filtered for depth (tumor sample coverage > 20x, normal sample coverage > 10x) and variant allele frequency (VAF ≥ 0.1). All mutations annotated to genomic regions not targeted by a sgRNA detected in at least one sample were kept.

### Identification of somatic copy number, structural variants and chromotripsis

Sequenza^51^ was used to derive somatic copy number profiles from WGS data using each sample’s matched normal sample. A modified version of the “copynumber” R package (https://github.com/aroneklund/copynumber) was used to create the necessary support objects and allowed for usage of the mm10 genome in the deployment of Sequenza on the mouse BAM files. Structural variations were called by Manta^52^ applying default settings. Chromotripsis events were called using ShatterSeek and following PCAWG guidelines^25^: high-confidence chromotripsis were called used ShatterSeek with visual inspection of number of structural variants > 6 and number of oscillating copy number > 6. Circos plots were generated with OmicCircos^53^.

### TCGA WGS and RNA data analysis

Chromothripsis score was calculated as follows: chromothripsis information of TCGA dataset were downloaded and summarized based on the label information for each chromosome then assign discete values: High confidence = 1; Linked to high confidence = 0.75; Low confidence = 0.5; Linked to low confidence = 0.25; No = 0. For each sample, a final value was calculated by multiplying the sum of these values by the number of chromosomes affected. EMT score was extrapolated from Gibbons and Creighton^32^.

### Slide-DNA-seq

Slide-DNA-seq was performed as previously described^22^. Bead arrays were produced and sequenced as described in Stickels et al, 2022. The bead oligonucleotide sequence is specific to slide-DNA-seq, and was produced in-house with the following nucleotide sequence:

5’-TTT_PC_GCCGGTAATACGACTCACTATAGGGCTACACGACGCTCTTCCGATCTJJJJJJJ JTCTTCAGCGTTCCCGAGAJJJJJJJNNNNNNNVVGCTCGGACACATGGGCG-3’

Where PC designates a photocleavable linker, J bases represent the spatial barcode (unique to each bead), N bases represent a unique molecular identifier (unique to each oligo), and V represents randomized bases that exclude T. Tissue handling, library preparation, and analysis were as described in Zhao et al., 2022, with the following modification. During the tagmentation stage, the Tn5 transposase solution was refreshed every hour, for a total of four rounds of tagmentation. All other experimental steps proceed exactly as previously described. Analysis steps were also performed as previously described, with scripts available at https://github.com/buenrostrolab/slide_dna_seq_analysis.

### scRNA-seq and GET-seq

scRNA-seq and scGET-seq were performed as previously described^28,54^. Briefly, scRNA-seq was performed on a Chromium platform (10x Genomics) using ‘Chromium Single Cell 3ʹ Reagent Kits v3’ kit manual version CG000183 Rev C (10x Genomics). Final libraries were loaded on a Novaseq6000 platform (Illumina) to obtain 50,000 reads *per* cell for scRNA-seq and 100,000 reads *per* nucleus for scGET-seq.

### Processing of scRNA-seq data

Cell barcodes were identified and quantified using UMI-tools^55^, setting the expected number of cells to 5,000. Read tags from 10x Chromium libraries were aligned to mm10 reference genome using STARsolo v2.7^56^. Count matrices were imported into scanpy objects^57^. Doublets were identified with scrublet^58^ using default parameters. After concatenation of all data samples, predicted doublets, cells with more than 20% counts assigned to mitochondrial genes and cells with less than 2,000 genes by counts were removed from the analysis. Counts were normalized and log1p transformed before cell cycle phase assignment. After that we calculated cell cycle progression value (CC_diff) as G2M_score – S_score. We regressed out CC_diff and %MT before scaling.

We calculated cell kNNgraph using BBKNN^59^ setting the number of neighbors to half the square root of the number of cells. Cell clusters were identified using schist^60^ with default parameters. CytoTrace score^61^ was calculated using cellrank^62^. Simpson and Shannon indeces were calculating on the frequency of cells belonging for each cluster in each group.

To compute the cell-to-cell transition matrix we applied a velocity kernel with dynamical scoring (vk) implemented in cellrank using the experimental time to set the progression. We also applied a connectivity kernel (ck) to stabilize the Schur decomposition. Finally, the transition matrix was calculated using a weighted kernel (0.8vk + 0.2ck). GPCCA method was then used to identify two major cell macrostates, assigned to mesenchymal or epithelial phenotypes. To quantificaty EMT gene signature we considered the difference of mean expression of genes up and down-regulated^63^. Activity of transcription factors (TF) was evaluated using decoupleR^64^ with default parameters.

### Processing of scGET-seq data

Raw reads were preprocessed as previously described^54^ using the scGET workflow^60^ with default parameters. Each sample consists of two matrices representing read counts for the mm10 genome binned at 5 kbp, one for Tn5 and one for TnH. Once count matrices have been generated, we fitted a zero-inflated Poisson distribution on Tn5 and TnH counts per each cell and generated the empirical distribution of the difference between the two transposases. Using such per-cell distribution, we computed the p-value of the difference Tn5-TnH for each genomic bin in each cell. p-values for each tail were log-transformed as previously detailed, so that accessible regions are assigned a positive value and compact regions are assigned a negative value.

We removed regions that were not profiled in at least 90% of the cells, we removed cells with a total coverage lower than 2,000. Reduction of dimensionality was achieved by tensor train decomposition (TTD) using tensorly python library. The first 20 components were used as basis to compute the cell kNN graph, using BBKNN^59^ with cosine distance (‘angular’) and number of neighbors equal to half the square root of the number of cells.

Chromatin Accessibility Score (CAS) was obtained as fraction of regions with enriched Tn5 over TnH, as results of the signed log(p-value) previously calculated.

We applied dot product of the matrix of signed p-values with the adjacency matrix of the kNN graph to obtain a smoothed matrix (Macc) of net accessibility values. This matrix was used to calculate a velocity kernel (vk) using cellrank. We regularized this kernel with a connectivity kernel (ck) (0.8vk + 0.2ck) before applying GPCCA method. Finally, two major cell macrostates were identified, assigned to mesenchymal or epithelial phenotypes.

To calculate the activity of TF from scGET-seq data we selected the top 200 genomic regions that could be considered drivers of each macrostate using cellrank correlation test. For each region we calculated the Total Binding Affinity^65^ of the whole set of transcription factors included in HOCOMOCO v11^66^.

To profile genome instability, we applied delly^67^ on aligned TnH reads to retrieve a list of structural variants (SV). As delly returns the name of reads supporting each SV, we matched read names to single cells to build a count matrix of SV. We applied a linear model fitting the log10-coverage (as number of TnH reads per cell) with the log(nSV + 1) (as number of SV per cell). We used the residuals of the fit as measure of genome instability. Positive values indicate an excess of SV at a given coverage (hence high instability).

To characterize the genetic heterogeneity in each group, we characterized Copy Number Alterations in single cells. To this end, we binned genome into 10 Mb windows and computed the per-cell coverage, then we fit ZIP on the coarse count matrix and calculated the hyperbolic arcsine of the ratio between counts and the expected number λ from the Poisson distribution, as a proxy of CNA values. The new matrix was processed do identify cell clusters using schist^60^. The size of intersections of such clusters with the macrostates was used to calculate ^2^D Hill numbers, α-diversity and β-diversity indices.

### Combined analysis of scGET-seq and scRNA-seq data

To integrate the scGET-seq and scRNA-seq data we applied MOWGAN, deep learning framework based on Generative Adversarial Network (WGAN-GP) that uses Wasserstein loss function and gradient penalty. The WGAN-GP training is performed in mini-batches, where cells are sampled from the whole dataset (*i.e.*, from each molecular layer). Input data for WGAN-GP are the single cell embeddings (PCA for scRNA-seq and TTD for scGET-seq), clipped to the same number of dimensions. Cells for each are sorted according to the first component of the Laplacian Eigenmap. Sampled data are combined in a tensor of shape (*N*, 2, *C*), where *N* is the number of cells in the mini-batch (256) and *C* is the number of components in each embedding used for the analysis. Mini-batches are sampled from the same experimental batches to avoid spurious associations. We select *N* cells from the scRNA-seq layer, then we select *n*=50 random mini-batches from scGET layer and run Bayesian Ridge regression to fit the best coupling. This step is repeated for every mini-batch that is used during training. The network generator is designed with three convolutional 1D layers and two batch normalization layers. The network critic is designed with two Conv1D layers and a Dense layer with 1 unit. All Conv1D layers are characterized by a kernel size of *M*, stride 1 and the ReLU activation function. Finally, different optimizers are used for each component: Adam optimizer for the generator with learning rate = 0.001, beta_1 = 0.5, beta_2 = 0.9, epsilon = 1*e* − 07 and the AMSgrad option; RMSprop optimizer with learning rate = 0.0005 for the critic. Once training has been performed, the MOWGAN can generate embeddings for synthetic cells that are artificially paired. For each modality, we created a single scanpy object for including the synthetic and the original data and applied Harmony^68^ to perform the integration. The new embeddings produced by Harmony were used to generate the kNN graphs for each modality. To identify cell states consistent across both modalities, we applied the multilevel Nested Stochastic Block Model, for which a single kNN is created with labeled edges corresponding to each modality.

### Statistics and Reproducibility

Data are presented as the mean or medians ± s.d and percentages. Comparisons among biological replicates were performed using two-tailed Student’s t test, two-way ANOVA, Mann-Whitney U test. Results from survival experiments were analyzed with log-rank (Mantel–Cox) test and expressed as Kaplan–Meier survival curves. Results from contingency tables were analyzed using two-tailed Fisher’s exact test or chi-test for multiple comparisons. All the statistical analysis have been performed with GraphPad Prism software. Data distribution was assumed to be normal without formal testing. Group size was determined on the basis of the results of preliminary experiments. No statistical methods were used to determine sample size. No data were excluded from the analysis. Group allocation and analysis of outcome were not performed in a blinded manner. *In vitro* experiments were repeated three times, while *in vivo* experiments were performed at least twice.

### Data availability

All data supporting the findings of this study are available within the Article and its Supplementary Information. Murine genomic and single cell RNA and GET seq raw data have been deposited on ArrayExpress under accession code: <u>E-MTAB-13051</u> and <u>E-MTAB-13052</u>. Previosly published datasets info are available within the supplementary information of this manuscript. Source data for *in vitro* and *in vivo* are provided with this paper. Requests for resources and reagent can be directed to the lead contact G.G.

### Code availability

Codes used for this manuscript have been previously published and adequately referenced in this manuscript. Methodological details on parameters used are available in the Methods section of this manuscript.

## Acknowledgments

We wish to thank all members of the Dr. Draetta and Viale lab for discussions and reagents. We would particularly like to thank Shan Jiang, and the MDACC Department of Veterinary Medicine, for valuable support in animals handling, Dr. Charles Kingsley, Jorge Delacerda, and the MDACC Small Animals Imaging Facility for their constant willingness. We thank Genscript and VectorBiolab for support and service. We wish to acknowledge the Advanced Technology Genomics Core (Erika Thompson, Dr. Hongli Tang, Steven Bates, David Pollock) at MDACC and the CA016672 (ATGC) Core Grant. The authors acknowledge the support of the High-Performance Computing for research facility at the University of Texas MD Anderson Cancer Center for providing computational resources that have contributed to the research results reported in this paper.

## Funding

F.Citron. was supported by AIRC and the European Union’s Horizon 2020 research and innovation program under the Marie Skłodowska-Curie grant agreement no. 800924. G.F.D. was supported by the Sheikh Ahmed Bin Zayed Al Nahyan Center for Pancreatic Cancer Grant and the Pancreatic Cancer Action Network Translational Research Grant. L.W. was supported by the NIH/NCI grants R01CA266280, the Cancer Prevention and Research Institute of Texas (CPRIT) awards RP200385, The Break Through Cancer (BTC) Foundation, the start-up research fund and the University Cancer Foundation via the Institutional Research Grant Program at the University of Texas MD Anderson Cancer Center. L.W. is also a Andrew Sabin Family Foundation Fellows at MD Anderson Cancer Center. G.G. was supported by the Barbara Massie Memorial Fund, the MDACC Moonshot FIT Program, the Bruce Krier Endowment Fund, and the Lyda Hill Foundation. K.L. is funded by the UK Medical Research Council (MR/P014712/1).

## Author contributions

Conceptualization: G.G., L.P., F.Chen, L.W., T.P.H., A.V., A.C., V.G., A.F., G.T., G.F.D., G.T., A.S.

Methodology: L.P., L.Z., S.M., A.R., F. Chen, D. Cittaro, G.G.

Software: L.P., L.Z., S.M., F.P., D.Cittaro

Formal analysis: L.P., L.Z., S.M., F.P., C.L.

Investigation: L.P., L.Z., S.M., F.G., F.P., F. Carbone, C.L., H.K., F. Citron, M.S., T.N.A.L., S.L., C.Z., D.C., E.G., N.F.

Visualization: L.P., L.Z., S.M.

Writing – Original draft: L.P., G.G.

Writing – Review and Editing: L.P., G.G., F. Chen, L.W., A.V.,A.C., G.F.D., A.S.

## Extended data Figures

**Extended data Figure 1.**
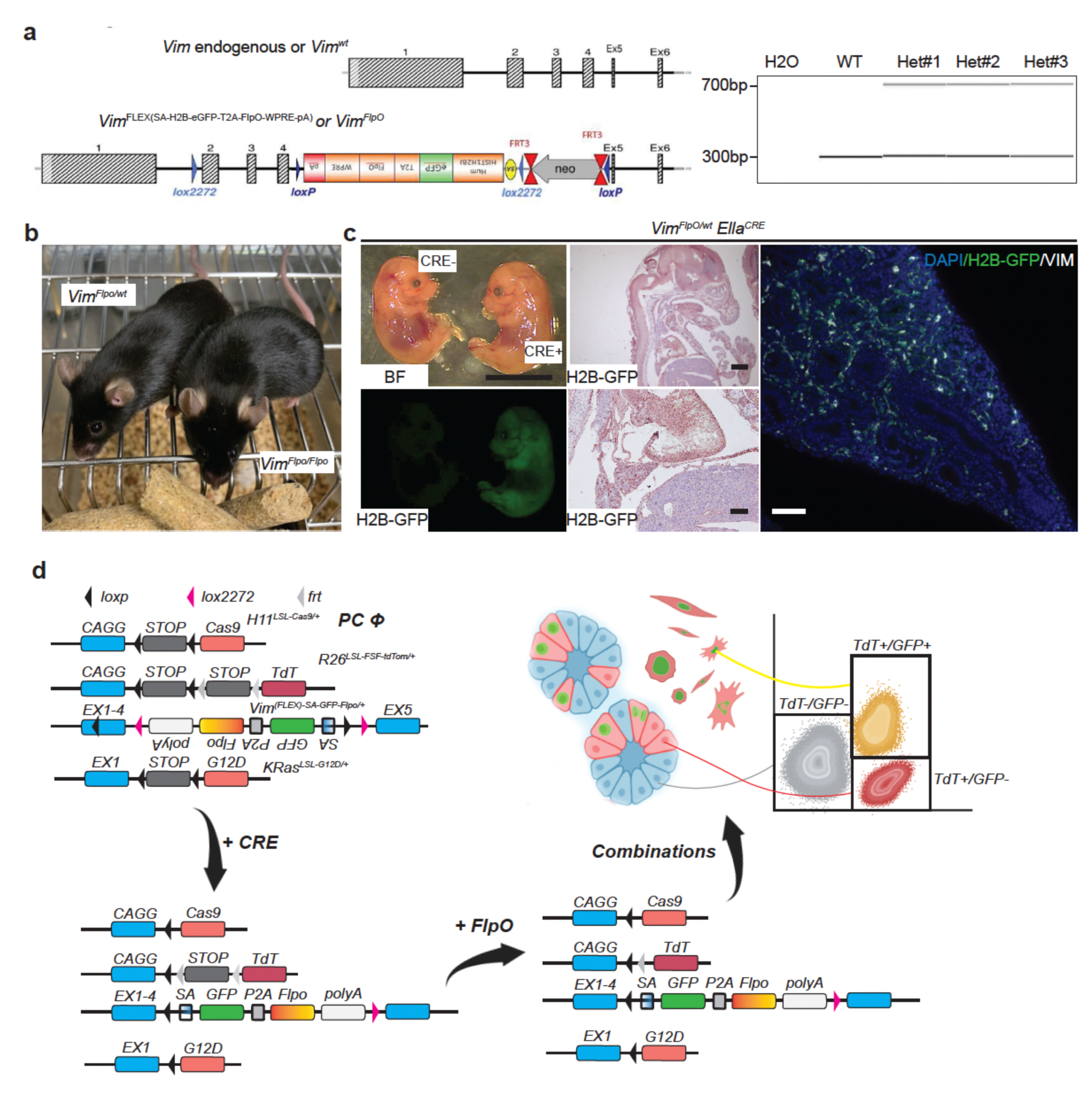
A SM-GEMM for lineage cells with mesenchymal features. **a-b)** Schematic showing strategy of the engineering of the mouse *Vimentin* (*Vim*) locus (*left*) and representative genotypes of heterozygous mice (*right*); mice were viable, fertile without any evident phenotype as shown by two adult heterozygous and homozygous littermates **(b)**. **c**) Representative macroscopic pictures showing detection of GFP fluorescence in *Ella^CRE^*, *Vim^FlpO/wt^* mice at embryonic day 13.5 (*left*) and histopathological examination of GFP and Vimentin (VIM) protein expression; it is readly noticeable the overlapping expression of GFP and VIM in developing lungs. Pictures are representative of N = 5 embryos. **d)** Schematic showing the SM-GEMM design and possible combinatorial outputs leading to a model able to lineage trace mesenchymal plasticity selectively in the tumor compartment.

**Extended data Figure 2.**
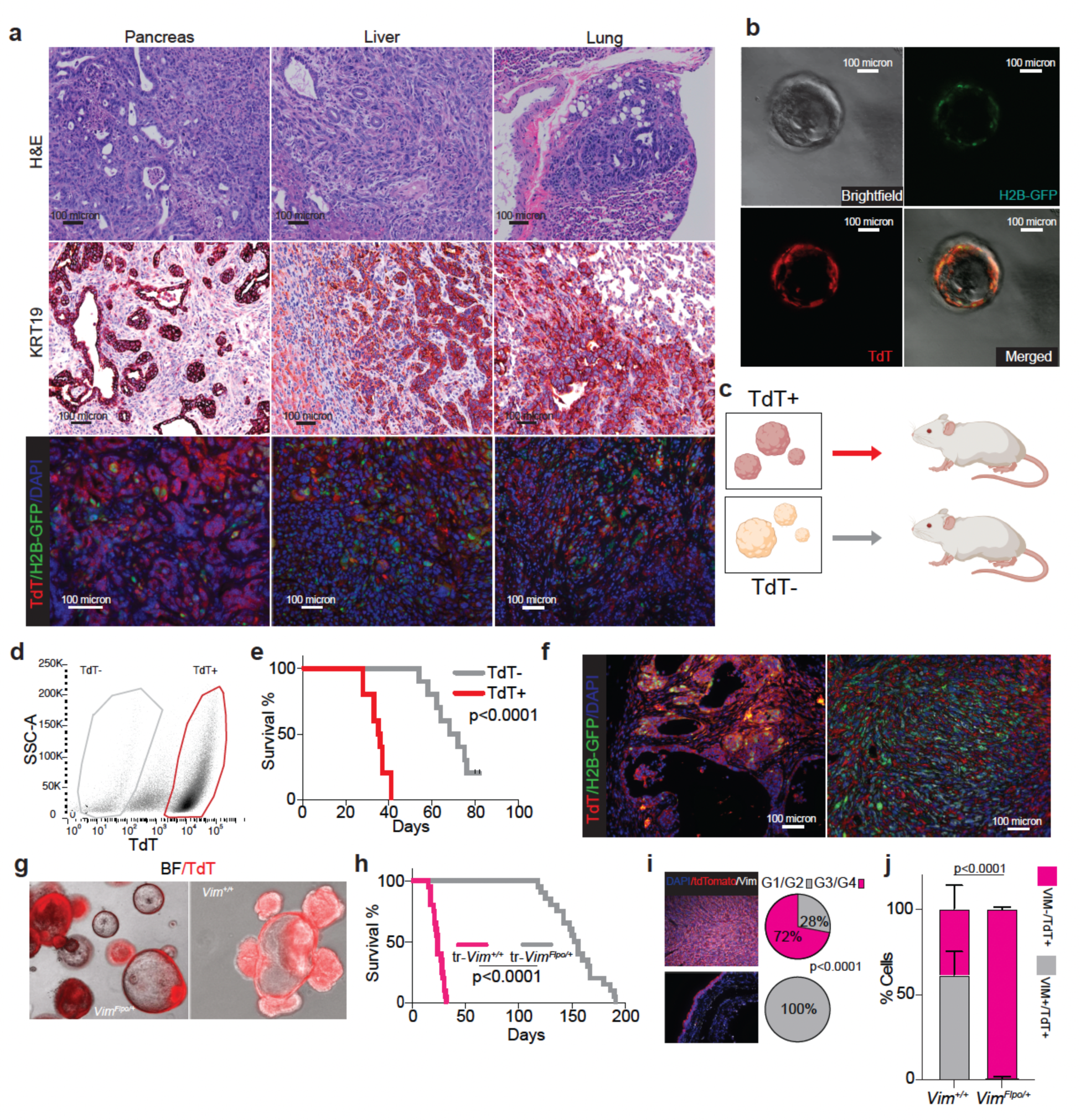
Pancreatic tumors with mesenchymal competency display aggressive clinical behavior. **a)** Histopathological characterization of primary tumor site and metastatic sites of PCΦ SM-GEMM showing distribution of TdT+/GFP^HIGH^ and TdT+/GFP^LOW^ cells in tumoral areas. **b)** Microscopical images of PCΦ SM-GEMM-derived organoids showing the concomitant presence of TdT+ and TdT-cells. **c-d)** Schematic of the isolation of TdT+ and TdT-cells from PCΦ SM-GEMM-derived organoids **(c)** and representative dot plot of gating strategies **(d)**. Dot plot is representative of N = 2 sorting experiments. **e-f)** Survival curves and representative histopathological analysis of the isolated TdT+ and TdT-cells sorted from PCΦ SM-GEMM-derived organoids showing reduced survival in mice bearing TdT+ derived lesions with tendency to sarcomatoid transformation; noticeably, TdT-cells derived tumors evolve to TdT+ lesions. N = 10 mice per group; data and images are representative of N = 2 experiments. **g-h)** Representative microscopic pictures of organoids derived from PCΨ *Vim^+/+^*and *Vim^Flpo/+^*SM-GEMM **(g)** and survival curves of these organoids transplanted into the pancreata of recipient mice **(h)**. N = 20 mice per group; data are representative of N = 2 experiments. **i)** Representative immunofluorescence (*left*) and grading (*right*) of tumors arose from transplanted organoids derived from PCΨ *Vim^+/+^* and *Vim^Flpo/+^*SM-GEMM. N = 25 tumor areas. **j)** Histopathological quantification of cells co-expressing VIM and TdT in tumors arose from transplanted organoids derived from PCΨ *Vim^+/+^*and *Vim^Flpo/+^*SM-GEMM. N = 25 tumor areas. P values are calculated as follows: **g,j)** Logrank Mentel-Cox test; **k)** Fisher’s exact chi test; **l)** two-way ANOVA.

**Extended data Figure 3.**
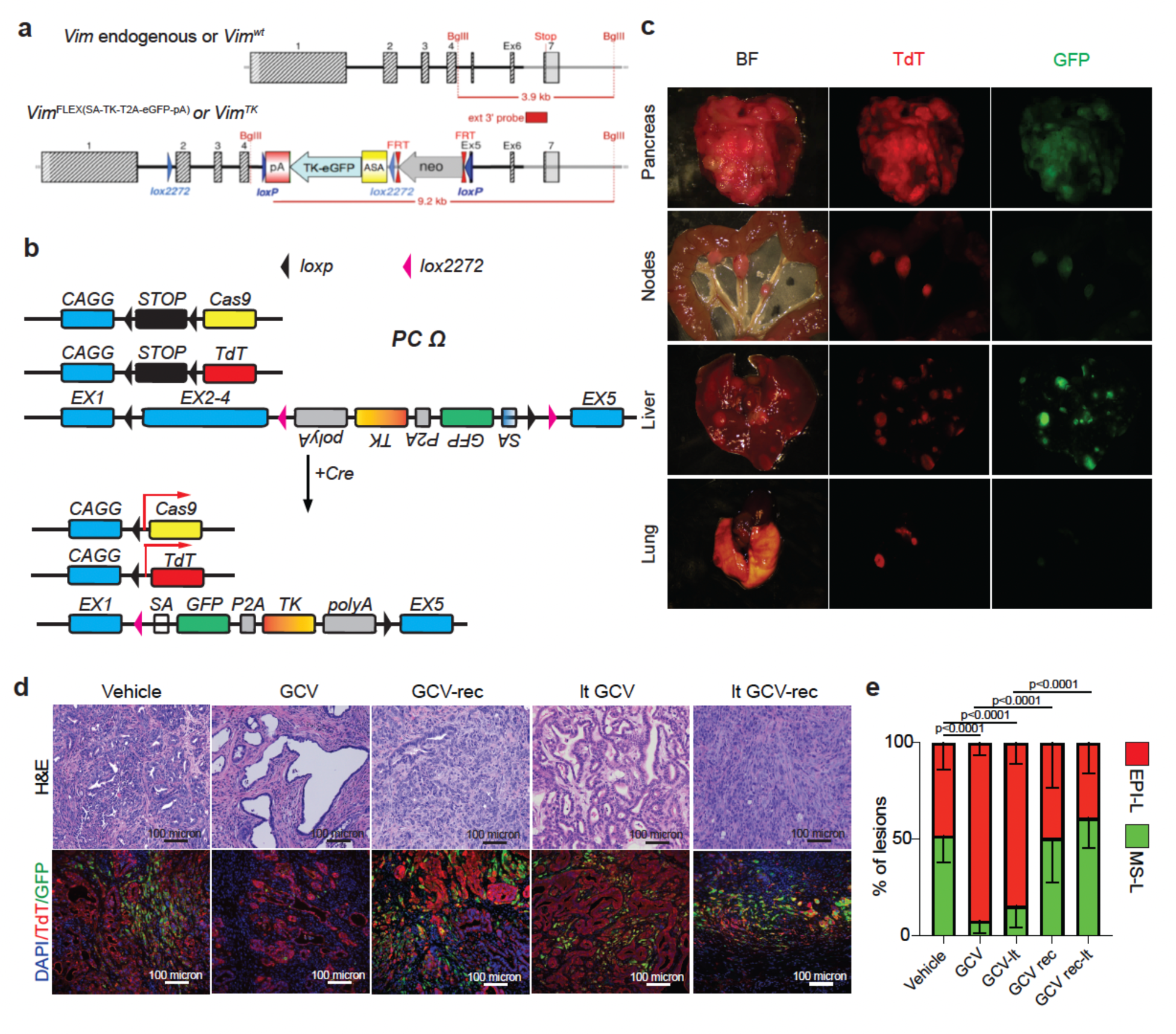
A SM-GEMM to selectively ablate cancer cells with mesenchymal competency. **a-b)** Schematics showing strategy of the engineering of the mouse *Vimentin* (*Vim*) locus with a eGFP-TK cassette for the selective ablation of cells expressing Vimenting upon GCV treatment **(a)** and outcomes after CRE mediated recombination **(b)**. **c)** Macroscopic pictures of primary tumors and metastatic lesions of a PCΩ SM-GEMM. Pictures representative of N = 20 mice. **d-e)** Histopathological analysis **(d)** and quantification **(e)** of primary tumor lesions for different treatment groups. It is noticeable the epithelial, ductual structures at the end of GCV treatment resulting in low grade, poorly aggressive phenotypes. P values are calculated as follows: **e)** two way ANOVA.

**Extended data Figure 4.**
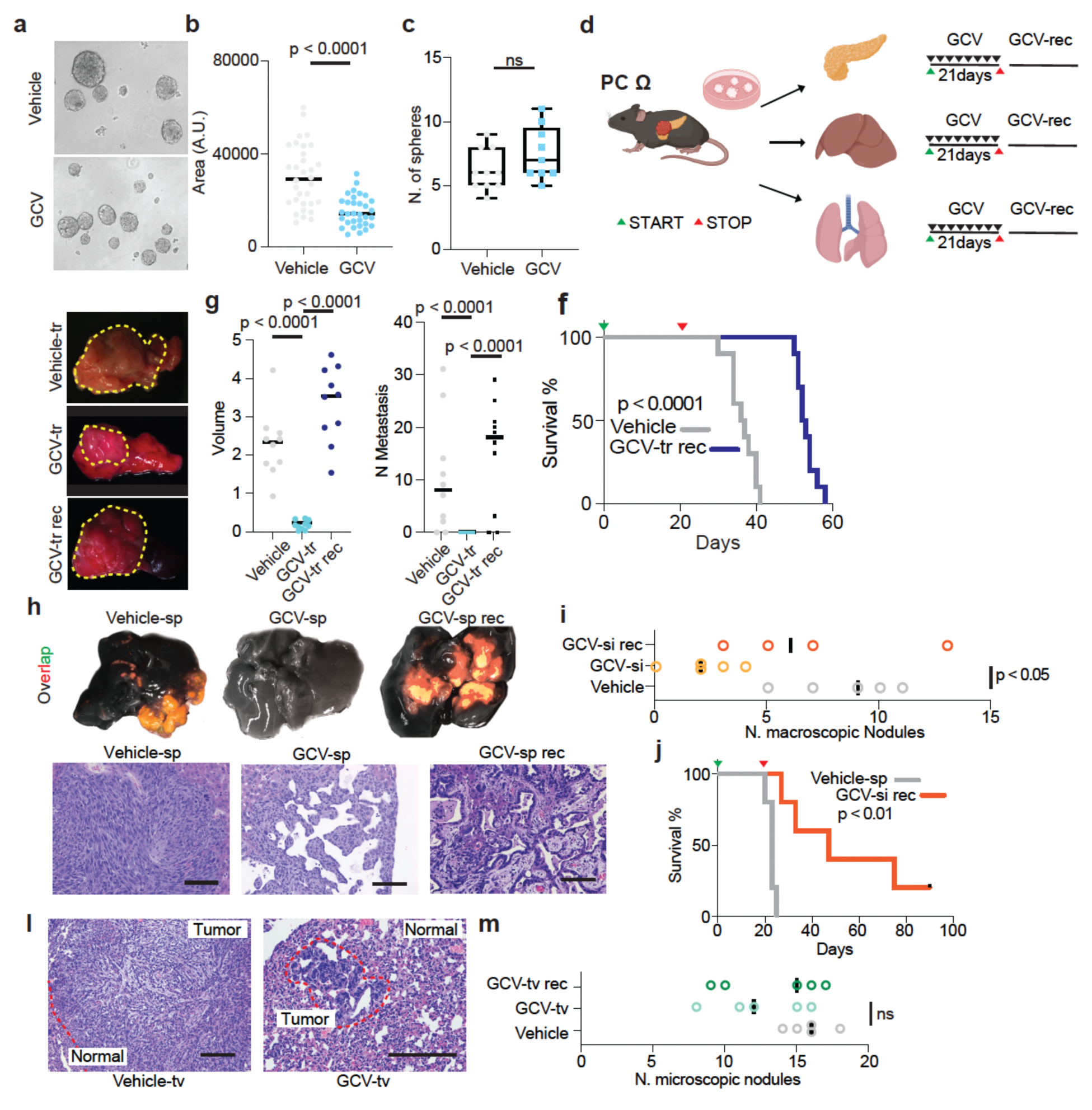
Cells with mesenchymal features are required for the expansion of tumor engraftments. **a)** Representative pictures of pancreatic tumor-derived 3D cultures generated from PCΩ SM-GEMM. Images representative of n = 3 independent primary cell lines. **b-c**) Dot plot with individual points (left) and box and whiskers plot showing area and number of pancreatic spheres upon GCV treatment. N = 32 fields per group (**b**) and N = 10 fields per group. **d)** Schematic showing transplantation experiments and treatment schedule of 3D cultures generated from PCΩ SM-GEMM. **e-f)** Representative macroscopic pictures of primary tumors for the three groups (left) and dot plots with individual values calculating tumor volume and number of metastasis for the same groups. Images are representative of 30 mice; N = 10 per group. **f)** Survival curves of mice bearing transplanted pancreatic tumor-derived 3D cultures upon orthotopic transplantation and GCV treatment. **h-i)** Representative macroscopic and microscopic pictures (left) and quantification of number of nodules (right) of pancreatic tumor-derived 3D cultures transplated via spleen injection and treated or not with GCV. Images are representative of N = 15 mice; N = 5 per group. **j)** Survival curves of mice bearing pancreatic tumor-derived 3D cultures upon splenic transplantation and GCV treatment. N = 5 per group. **l-m**) Representative histopathological pictures (left) and number of nodules (right) from mice transplated in the lateral vein with pancreatic tumor-derived 3D cultures. Images are representative of N = 15 mice; N = 5 mice per group. P values are calculated as follows: **b,c,f,i,m)** two-sided student t-test; **g, j)** Logrank Mentel-Cox test.

**Extended data Figure 5.**
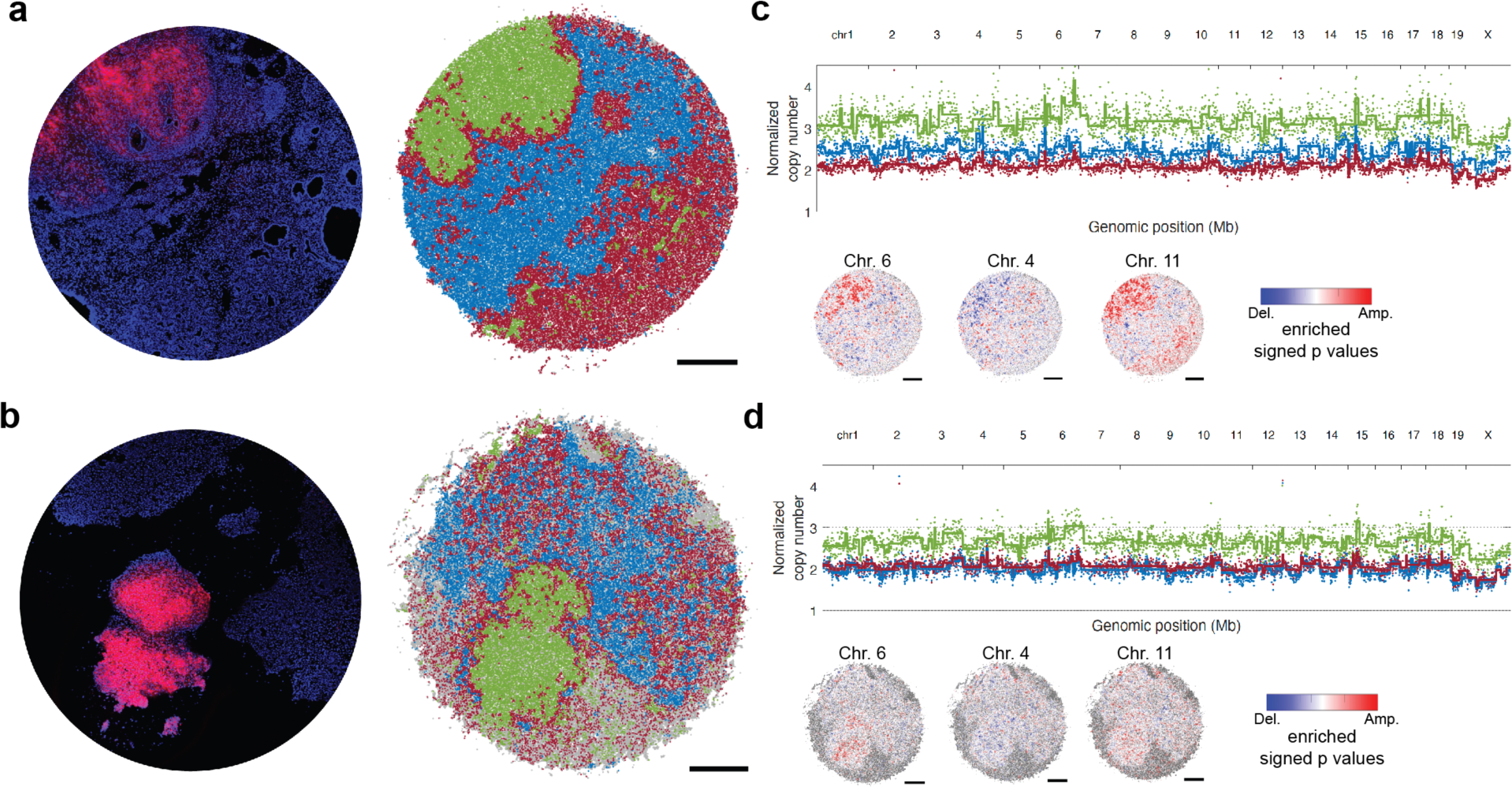
Spatial genomics of tumor cells with mesenchymal proficiency. **a-b)** Slide-DNA-seq of two PCΦ SM-GEMM 3 weeks after AAV transduction showing TdT+ and TdT-regions of pancreatic tissue, differentially clustering according to genomic analysis. **c-d)** Normalized copy number scatter plots showing differential copy number aberrations in the three clusters as previously calculated with principal component analysis; particularly evident are gains of 6 and 15 and minor loss of chromosome 4.

**Extended data Figure 6.**
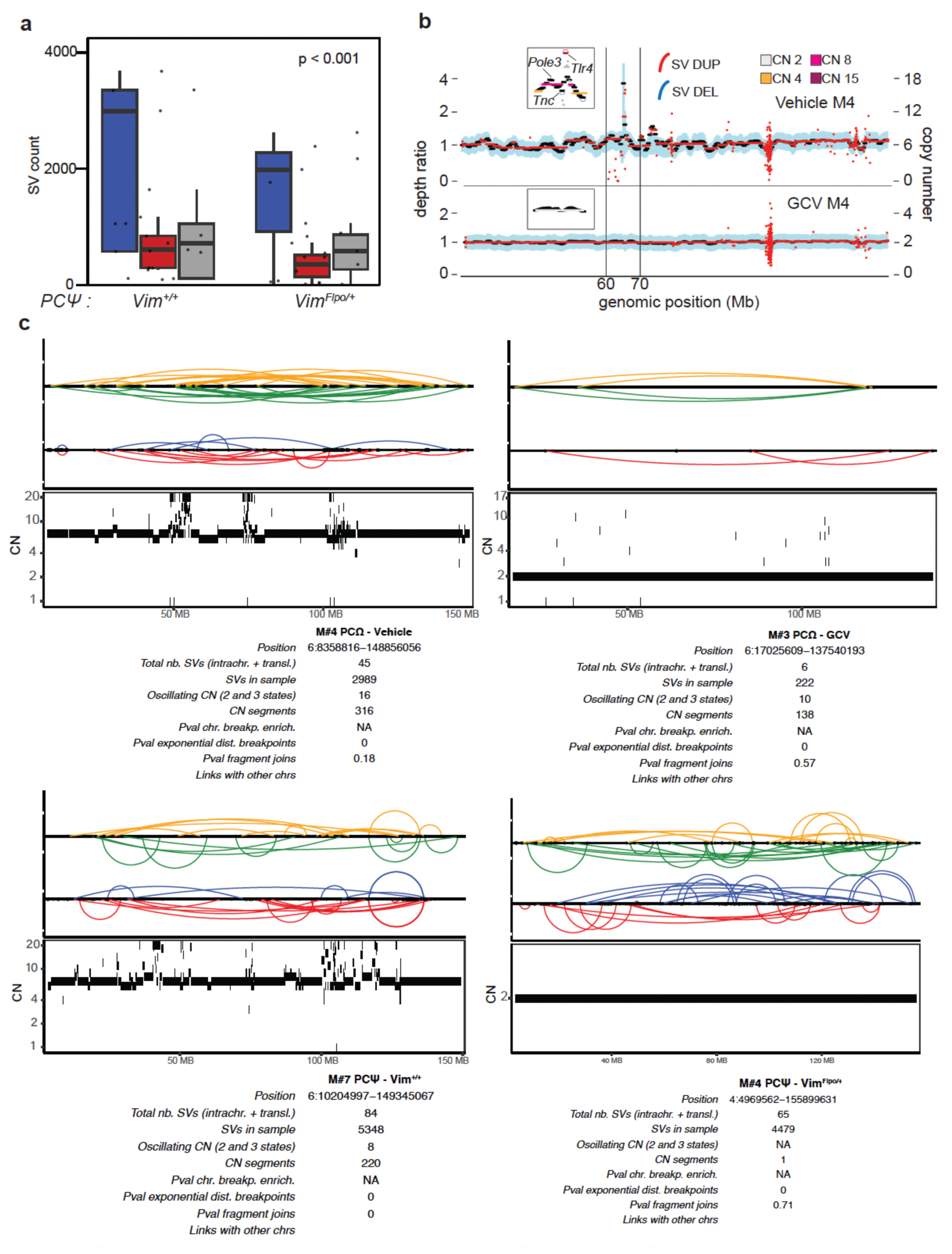
Structural variants and chromotripsis in EMT-on and EMT-off groups. **a)** Box and whiskers with individual points showing number and type of SVs in the PCΨ Vim^+/+^ and Vim^Flpo/+^ SM-GEMMs. N = 7 mice per group. **b)** Two representative scatter plots showing absolute copy number values of chromosome 4 for two different mice belonging to the EMT-on (top) and EMT-off (bottom groups). Poliploidy and focal copy number gains can be readily appreciated. **c)** ShatterSeek output plots for 4 different mice belonging to the EMT-on (left) and EMT-off (right) group. Visual inspection, SVs count and number of oscillating copy number (Oscillating CN) were used to call high-confidence chromotripsis: the two cases on the left met the criteria (**Methods**).

**Extended data Figure 7.**
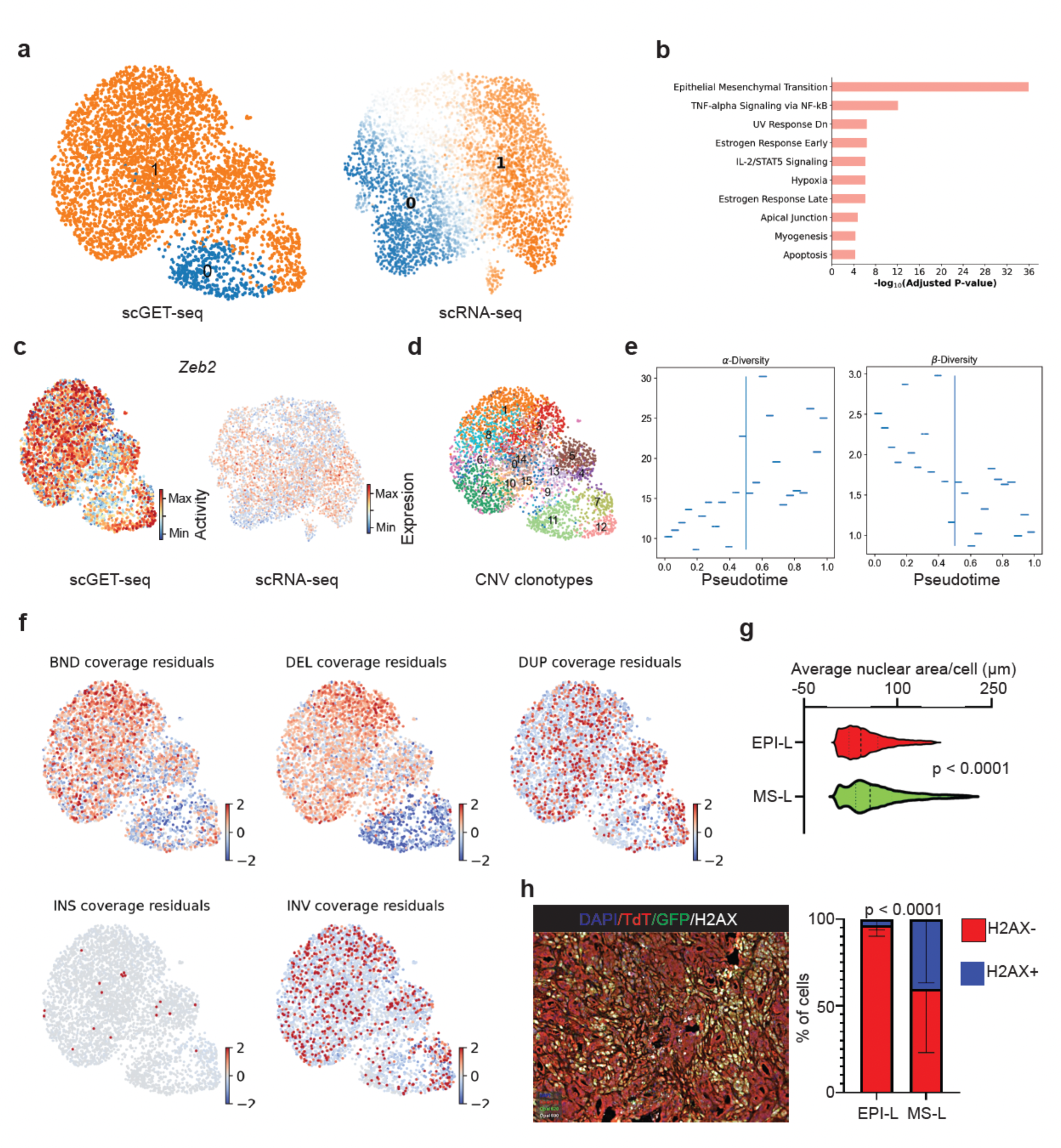
Genomic instability and genetic heterogeneity correlates with transcriptomic and epigenetic features of EMT. **a-b)** UMAPs showing cluster distribution of MS-L and EPI-L tumor cells and top 10 significant pathways enriched in cluster n.1 according to scGET and RNA data integration. **c)** UMAPs showing the distribution of *Zeb2* EMT-related transcription factors enriched in cluster n.1. **d)** UMAP showing clonotype distribution as calculated by copy number analysis of scGET seq data. **e)** Scatter plot displaying correlations between Pseudotime α-Diversity and β-diversity as calculated from scGET seq data. **f)** UMAP showing distribution of structural variants as calculated from scGET seq data after normalization of sequencing coverage. **g)** Violin plot showing average nuclear area per cell as calculated by histopathological analysis of PCΩ SM-GEMMs. **h)** Histopathological representative picture (left) and quantification (right) of PCΩ SM-GEMMs with the DNA damage marker γH2AX. P values are calculated as follows: **g)** two-sided student t-test; **h)** two-way ANOVA.

**Extended data Figure 8.**
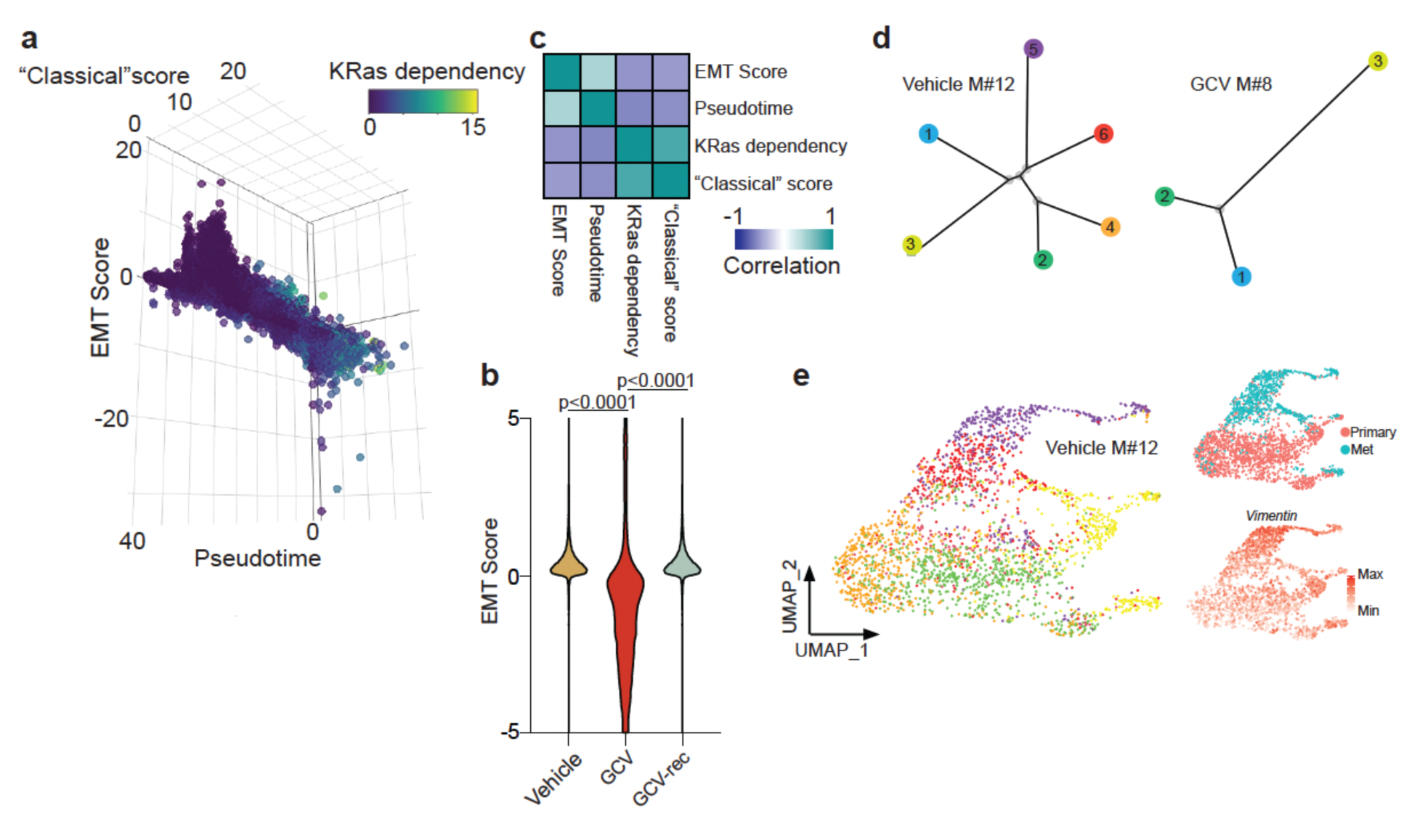
EMT unlocks transcriptomic heterogeneity in PDAC. **a)** Three dimensional dot plot showing distribution of cells along the Pseudotime, EMT Score and “Classical” score (**Methods**); it is noticeble that cells with mesenchymal transcriptomic features are indipendent from the oncogene signature. **b)** Violin plot showing EMT Score values for the three groups included in the scRNA seq analysis. **d)** Inferred trees of clonotypes as calculated by SCEVAN algorithm (**Methods**, **d**) for two representative cases of PCΩ Vehicle and GCV groups. **e)** UMAP showing clonotype distribution in primary and metastatic lesions together with *Vimentin* gene expression. P values are calculated as follows: **b)** two-sided Mann-Whitney test.

## Notes

### Competing Interest Statement

The authors have declared no competing interest.

